# Biological convolutions improve DNN robustness to noise and generalisation

**DOI:** 10.1101/2021.02.18.431827

**Authors:** Benjamin D. Evans, Gaurav Malhotra, Jeffrey S. Bowers

## Abstract

Deep Convolutional Neural Networks (DNNs) have achieved superhuman accuracy on standard image classification benchmarks. Their success has reignited significant interest in their use as models of the primate visual system, bolstered by claims of their architectural and representational similarities. However, closer scrutiny of these models suggests that they rely on various forms of shortcut learning to achieve their impressive performance, such as using texture rather than shape information. Such superficial solutions to image recognition have been shown to make DNNs brittle in the face of more challenging tests such as noise-perturbed or out-of-domain images, casting doubt on their similarity to their biological counterparts. In the present work, we demonstrate that adding fixed biological filter banks, in particular banks of Gabor filters, helps to constrain the networks to avoid reliance on shortcuts, making them develop more structured internal representations and more tolerant to noise. Importantly, they also gained around 20−35% improved accuracy when generalising to our novel out-of-domain test image sets over standard end-to-end trained architectures. We take these findings to suggest that these properties of the primate visual system should be incorporated into DNNs to make them more able to cope with real-world vision and better capture some of the more impressive aspects of human visual perception such as generalisation.

## 1 Introduction

The success enjoyed by deep convolutional neural networks (DNNs) in complex perceptual tasks, notably image classification, has led many researchers to suggest that they accomplish their objectives in a similar manner to humans. Architectural and representational similarities further reinforce this view of DNNs, not just as engineering tools, but as good models of primate vision (Cadena et al., 2019; Guclu & van Gerven, 2015; Khaligh-Razavi & Kriegeskorte, 2014; Kubilius et al., 2016; Kubilius et al., 2018; Schrimpf et al., 2018; Yamins et al., 2014; Yamins & DiCarlo, 2016). However, in stark contrast to humans, one of the most striking failures of these models is their lack of ability to generalise outside of their training sets. This casts doubt on the claims that such models work in a fundamentally similar way to humans.

In contradiction to earlier claims that DNNs learn about object shape as a representational basis for their image classifications (Kriegeskorte, 2015; Kubilius et al., 2016; LeCun et al., 2015), subsequent work has found a strong bias towards textures and similar spatially high-frequency information (Baker et al., 2018; Deza & Konkle, 2021; Geirhos et al., 2019). Likewise in our earlier work, we reported that in the extreme, standard DNNs would base their image classifications on just a single pixel when correlated with image category, disregarding the richer shape information (Malhotra et al., 2020).

The tendency of DNNs to solve tasks in unintended ways has been characterised as “shortcut learning”, whereby decision rules are learnt which facilitate high performance on standard benchmarks but fail to generalise to more challenging test sets (Geirhos, Jacobsen et al., 2020). In this vein, a range of weaknesses of DNNs have been identified, including susceptibility to adversarial attacks (Szegedy et al., 2014), bias amplification (Bolukbasi et al., 2016) and intolerance to noise (Geirhos et al., 2018). Similarly, other authors have characterised these shortcomings as the models learning to rely on “non-robust” features that are present in the training data (Ilyas et al., 2019). While these problems could be regarded as properties of the dataset which fail to capture the richness of the visual world, we argue that they stem from insufficient *inductive biases* constraining the model to find more robust and general solutions. To frame it more positively, robust generalisation needs good inductive biases (Feinman & Lake, 2018; Lake et al., 2017; Sinz et al., 2019).

Inductive biases may be incorporated into the three core components of artificial neural network design: the objective function, the learning rule and the architecture (Richards et al., 2019), in addition to the training data (“environment”). In the present work, we focus on architectural constraints in the form of prescribed kernels in the first convolutional layer(s), taking inspiration from the receptive fields found in the early primate visual system. This particular form of inductive bias has received relatively little attention in the deep learning community, with a strong preference to instead rely upon full end-to-end training. This is a stark departure from the hand-tuned, featuring-engineering approach of classical computer vision research, despite some of this early work being encouragingly biologically plausible (Akbarinia & Parraga, 2018a, 2018b; Alahi et al., 2012; Riesenhuber & Poggio, 1999).

Although this approach has led to state-of-the-art scores on common benchmarks, end-to-end trained artificial neural networks (ANNs) have nonspecific (weak) biases and learn the statistics of the training data which may not generalise to out-of-distribution (*o*.*o*.*d*.) data (Sinz et al., 2019). Indeed, recent work on contrast normalisation demonstrated that even a slight deviation from the training distribution is enough to trigger a failure to generalise (Akbarinia & Gil-Rodísiguez, 2020). Arguably, this has become to an example of Goodhart’s Law (Strathern, 1997), where DNNs further surpass human performance on common image recognition benchmarks, yet no longer represent good measures as they fail to capture many interesting and elementary properties of visual perception.

While end-to-end training typically yields features resembling Gabor filters, an array of other filters emerge which lack a clear correspondence to those observed in the early visual system, further suggesting that DNNs are under-constrained (Krizhevsky et al., 2012, Fig. 3). As expected from the “bias-variance tradeoff” in supervised learning, the approach of fixing early convolutional forms has not (yet) achieved such high performance scores on standard benchmarks as with full end-to-end training. However, our previous results suggest that they may encourage DNNs to develop more robust and generalisable representations (Malhotra et al., 2020; Malhotra et al., 2019).

Furthermore, there is a strong motivation to fix the early convolutions from both the perspective of natural image statistics (Bell & Sejnowski, 1997; Olshausen & Field, 1996) and a developmental biology perspective (Briggman et al., 2011). Useful motifs about stable properties of the environment are most likely to pass through the “genomic bottleneck” conferring an evolutionary advantage by alleviating the burden on the individual to learn them (Zador, 2019), especially if they are “perceptual universals” of the world (Shepard, 1994).

Early work with DNNs showed how kernels strongly resembling Gabor filters naturally arise through training on naturalistic images (along with more obscure filters) (Krizhevsky et al., 2012) while recent computational modelling has even demonstrated how the particular hierarchy of receptive fields may arise from the retinal bottleneck (Lindsey et al., 2019). If centre-surround and Gabor filters form a visual alphabet of the natural world then they should be pre-wired (Gaier & Ha, 2019) or fixed rapidly due to evolution-optimised architectures (Zador, 2019) and remain relatively stable throughout the lifetime of the individual (and so also in models). In contrast to classical computer vision approaches, the features of the early layers are not “*hand*-engineered”, but essentially “*evolution*-engineered”.

Besides potential gains in “real-world” use (through increased resilience to noise and better *o*.*o*.*d*. generalisation), constraining DNNs with biologically-inspired inductive biases may also help to make them more interpretable by encouraging them to develop internal representations which are better aligned with their biological counterparts. This would potentially be a useful development for shining a light on otherwise obscure “black-box” models, allowing their decision processes to be better understood, refined, and overridden when necessary. Accordingly, we examine the most activating features of the trained models to visualise the differences in their internal representations.

Early work with Gabor kernels in convolutional neural networks focussed on the energy efficiency gains and speed of training convergence afforded by having fewer modifiable parameters while maintaining a structure conducive to image classification (Alekseev & Bobe, 2019; Meng et al., 2019; Sarwar et al., 2017). However, like other promising research with biologically motivated front-ends, without further constraining the parameters of the Gabor kernels, the models develop an over-reliance on the spatially high-frequency filters and forfeit their robustness to noise (Wu et al., 2019).

In our previous work with Gabor-kernel convolutions, the filters acted as a kind of regulariser, steering the network away from relying upon non-robust (yet diagnostic) features towards more robust representations (Malhotra et al., 2020; Malhotra et al., 2019). Subsequent work using *>*20–40× more Gabor filters demonstrated more resilience to adversarial attacks and noise perturbations over the corresponding end-to-end trained models (Dapello et al., 2020). Their study showed that the single biggest factor in attaining this improvement was the inclusion of stochasticity (Gaussian noise), particularly during training. This further suggests that the modifications worked to help the model develop more robust representations, in a way accounted for in earlier work by training on similar noise to the test set (Geirhos, Temme et al., 2020).

In the work presented here, we specifically examined the form of fixed kernels in the early convolutional layers of otherwise standard DNNs for their effects on internal representations, robustness to noise, and generalisation beyond the training set. In particular, we investigated a very human *o*.*o*.*d*. generalisation ability — to classify images based on simple line drawings (Hochberg & Brooks, 1962), their global shape features or their bounding contours rather than local textures (Baker et al., 2018).

We hypothesised that biologically inspired filter banks would make the models (a) more robust to noise perturbations applied to *i*.*i*.*d*. images, (b) better able to generalise to *o*.*o*.*d*. images and (c) develop more interpretable internal representations. Our results support these hypotheses for several types of common noise perturbations, reveal a 20 − 35% improvement in accuracy on our novel generalisation test sets and demonstrate striking differences in the internal representations.

## 2 Methods

Standard deep convolutional neural networks were trained with full end-to-end learning to obtain their baseline performance on image classification tasks. Each model architecture was then modified by configuring the first convolutional layer(s) to have fixed banks of kernels for each of several forms described below. These modified models were then trained on the same images as the standard models for 100 epochs to ensure that they reached convergence. The models were then compared by their performance on noise-perturbed validation images, generalisation test images and their internal representations. The models were implemented with Keras and Tensorflow 2. All simulation and analysis code (written in Python 3) is open-source and available at github.com/bdevans/BioNet.

### 2.1 Models

Several standard DNN architectures were used including `ALL-CNN` (Springenberg et al., 2015), `ResNet50` (He et al., 2016) and `VGG-16` (Simonyan & Zisserman, 2015). For each model, either the original architecture was used (“Original”) for full “end-to-end” training or the first convolutional layer was replaced with a bank of unmodifiable kernels. These fixed kernels took one of the following specific forms: Gabor, Difference of Gaussians (DoG) or Low-pass filters (chosen as a non-biologically motivated alternative way to smooth out noise). A “Combined” front-end was also used, whereby the first convolutional layer of a standard DNN was replaced with two fixed convolutional layers consisting of a DoG layer followed by a Gabor layer, modelling the receptive field organisation of the early visual stream. Each fixed kernel was set to 63 × 63 pixels in order to allow the filters to be adequately expressed without significant truncation at the edges, over a biologically relevant range of spatial scales. In the case of the Combined front-end, the kernels were reduced to 31 × 31 pixels due to computational constraints. The choice of (other) parameters for these convolutional kernels are given in Table 1 and the resulting kernels are visualised in Figure 1.

**Table 1:**
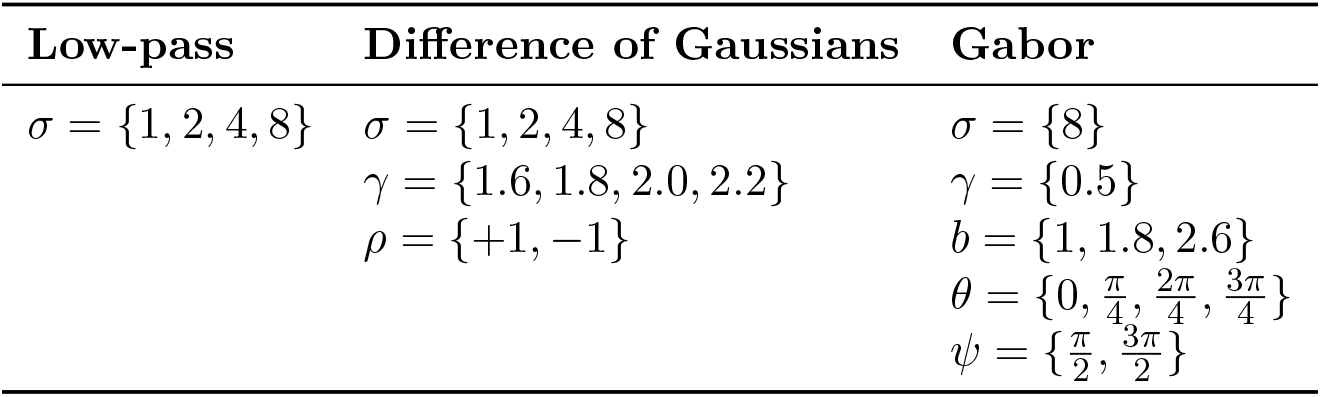
Parameters of the fixed convolutional kernels.

**Figure 1:**
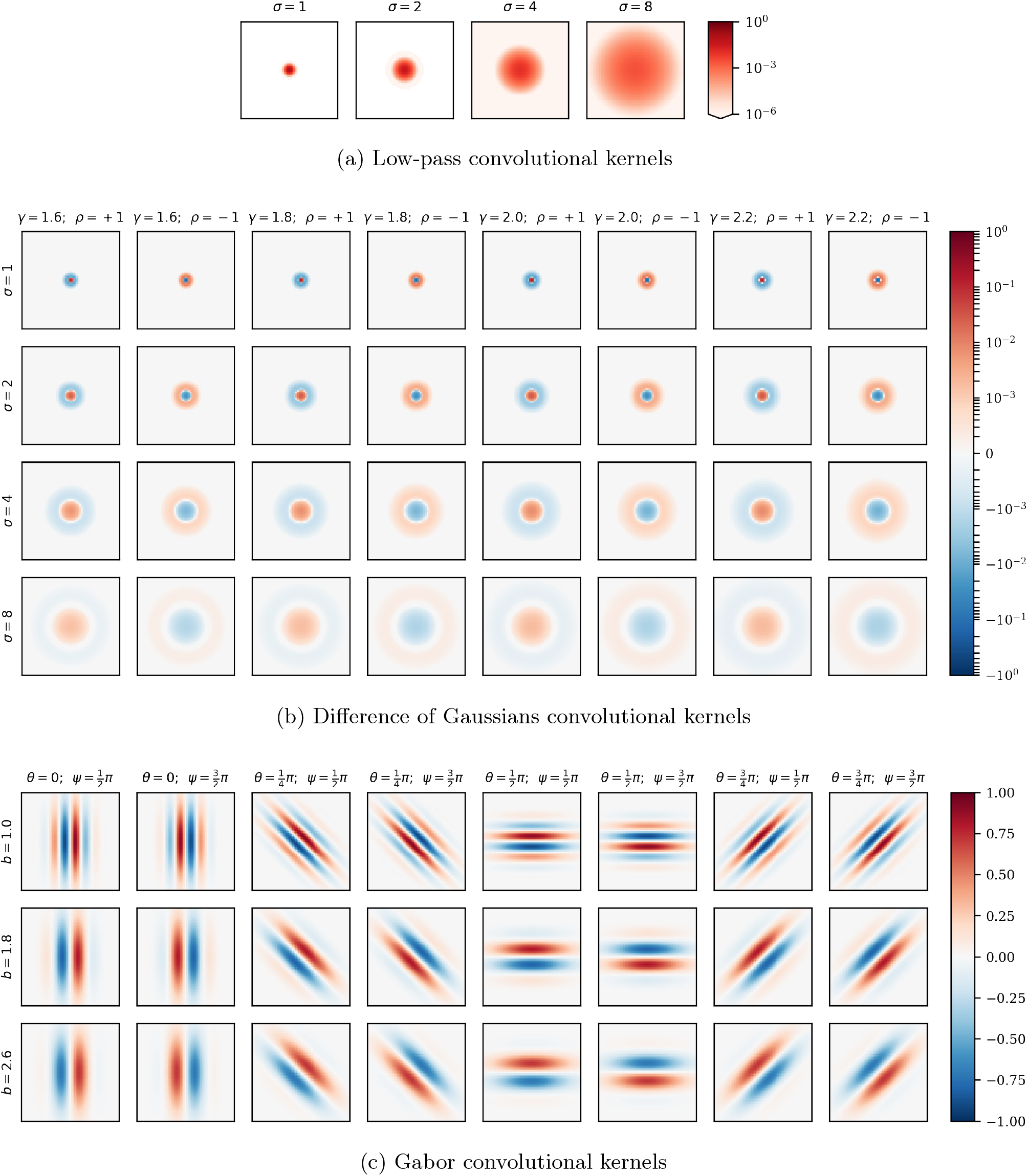
Illustration of the banks of fixed kernels used in the first convolutional layer(s).

In all cases, the input layer was modified to reflect the upscaled image size and conversion to greyscale, leaving only one luminance channel (224 × 224 × 1) as described in Section 2.3. Similarly, the output layer was reduced to classify each images into one of the 10 categories of `CIFAR-10`.

#### 2.1.1 Fixed convolutional kernels

##### Low-Pass

Low-pass filters were implemented as a simple 2-dimensional Gaussian kernel (Equation 1) which was convolved with the inputs, effectively blurring them by a degree parameterised by *σ*, the standard deviation of the Gaussian.

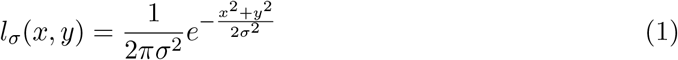

In the models presented, four channels (corresponding to four values of sigma) were used for the low-pass front-end as detailed in Table 1 and are shown in Figure 1a.

##### Difference of Gaussian

The Difference of Gaussians kernel (Equation 2) is the result of a surround Gaussian subtracted from a (typically smaller) centre Gaussian. The standard deviation of the centre Gaussian is parameterised by *σ* and the standard deviation of the surround Gaussian is parameterised by *γ* · *σ* where *γ* ≥ 1. In this work, the difference in Gaussians is multiplied by *ρ* ∈ {+1, −1} to model “on-” and “off-centre” ganglion cell receptive fields respectively.

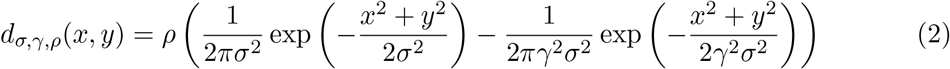

The Difference of Gaussians front-end had a total of 32 channels from combining values for three parameters as described in Table 1 and are shown in Figure 1b.

##### Gabor

The Gabor function is an oriented sinusoidal grating convolved with a Gaussian envelope (Equations 3-5) where *x* and *y* specify the position of a light impulse in the visual field (Petkov & Kruizinga, 1997).

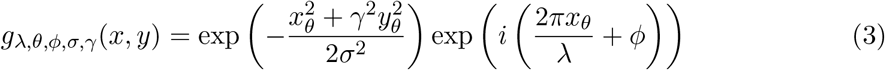

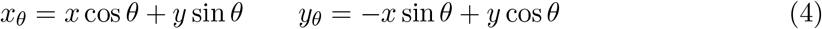

Rather than specify the wavelength of the sinusoidal component (*λ*) in pixels, it is more natural to set the bandwidth, *b*, which describes the number of cycles of the sinusoid within the Gaussian envelope, (which has a fixed standard deviation, *σ*, matched to the other front-ends). The wavelength of the sinusoidal factor, *λ*, is therefore set indirectly through *b*, and *σ* :

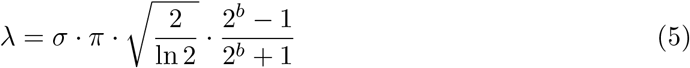

The Gabor front-end used had a total of 24 channels from combinations of values across its five parameters, chosen to span a range matched to primate primary visual cortex (Petkov & Kruizinga, 1997), shown in Table 1 and visualised in Figure 1c.

In the case of the Combined front-end models, the kernels of the first two convolutional layers are as shown in the DoG and Gabor plots (Figure 1b&c), however the kernel (canvas) size was reduced to 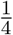 of their size (31 × 31 pixels) due to memory limitations.

### 2.2 Training

All models were trained with the modified (224 × 224 and greyscale) `CIFAR-10` training images (unperturbed and shuffled) to minimise categorical cross-entropy using Stochastic Gradient Descent (SGD) with a batch size of 64, a learning rate of 10^−4,^ and a decay of 10^−6,^. Training proceeded for 100 epochs, reducing the learning rate on plateau (after 5 epochs) by a factor of 0.2. Each model architecture was trained for five different random seed initialisations (eliminating seeds which failed to train) on NVIDIA GPUs.

Example kernels learnt in the first convolutional layer through this training procedure are illustrated in Figure 11. These 64 kernels are taken from a `VGG-16` model (modified and trained as described) with each being 3 × 3 pixels in size.

### 2.3 Stimuli

In all cases, the training images were based on the `CIFAR-10` dataset (which contains 10 classes of 6, 000 images per class, with 1, 000 of each held out for validation, see www.cs.toronto.edu/∼kriz/cifar.html). For testing, three categories of images were used; `CIFAR-10` validation images, noise-perturbed `CIFAR-10` validation images or generalisation image sets (described later).

To simplify the filter banks, we converted all images to greyscale according to the ITU BT.601 luma transform conversion formula (*Y* = 0.299·*R*+0.587·*G*+0.114·*B*), which models the trichromatic sensitivities of the human eye. Using a method similar to (Geirhos et al., 2018), the `CIFAR-10` images were then upscaled from their original dimensions of 32 × 32 pixels to 224 × 224 pixels using Lanczos resampling with luminosities clipped to [0, 255]. Each image was further preprocessed before presentation to the network by rescaling the intensity values from [0, 255] to [0, 1]. Under testing conditions where the images were perturbed, noise was applied after this rescaling, then the values were clipped in the range [0, 1] before rescaling back to the range [0, 255], as expected by the standard DNN architectures.

The mean and standard deviation were calculated across the entire (modified) training set and used for feature-wise centring and normalisation. Data augmentation was used to randomly shift the images vertically and horizontally by up to 10% (24 pixels) and to randomly apply a horizontal flip.

#### 2.3.1 Noise perturbations

Building on the work of (Geirhos, Temme et al., 2020) we explored the robustness of representations developed in DNNs with the range of different trainable and fixed convolutional kernels described. The `CIFAR-10` validation images were perturbed with a battery of common types of noise, systematically spanning a range of severity, before being presented to the networks. A summary of these noise perturbations is given in Table 2 with an illustration of them applied to one of the validation images in Figure 2.

**Table 2:** Image perturbation descriptions and severity.

| Perturbation | Description | Levels |
| --- | --- | --- |
| Uniform | Pixel-wise additive uniform noise drawn from $[-w, +w]$ then clipped at $[0, 1]$. | $w \in \{0, 0.1, \dots, 0.9, 1.0\}$ |
| Salt and Pepper | Pixels are randomly set to either black or white with probability, $p$. | $p \in \{0, 0.1, \dots, 0.9, 1.0\}$ |
| High-Pass | High-pass filtering with standard deviation of the Gaussian filter, $\sigma$. | $\sigma \in 10^{\{2, 1.8, \dots, 0.2, 0\}}$ |
| Low-Pass | Low-pass filtering with standard deviation of the Gaussian filter, $\sigma$. | $\sigma \in 10^{\{0, 0.2, \dots, 1.8, 2\}}$ |
| Contrast | Contrast, $c$ adjusted by setting each pixel intensity, $i$, according to $i' = (1 - c)/2 + i \cdot c$. | $c \in \{1, 0.9, \dots, 0.1, 0\}$ |
| Phase Scrambling | Phases are randomly shifted (in the Fourier domain) in the interval $[-w, +w]$ degrees. | $w \in \{0, 18, \dots, 162, 180\}$ |
| Darken | Each pixel intensity, $i$, is reduced by $l$. | $l \in \{0, 0.1, \dots, 0.9, 1\}$ |
| Brighten | Each pixel intensity, $i$, is increased by $l$. | $l \in \{0, 0.1, \dots, 0.9, 1\}$ |
| Rotation | Each image is rotated by $\theta$ degrees. | $\theta \in \{0, 90, 180, 270\}$ |
| Inversion | Pixel intensities are inverted. | $v \in \{0, 1\}$ |

**Figure 2:**
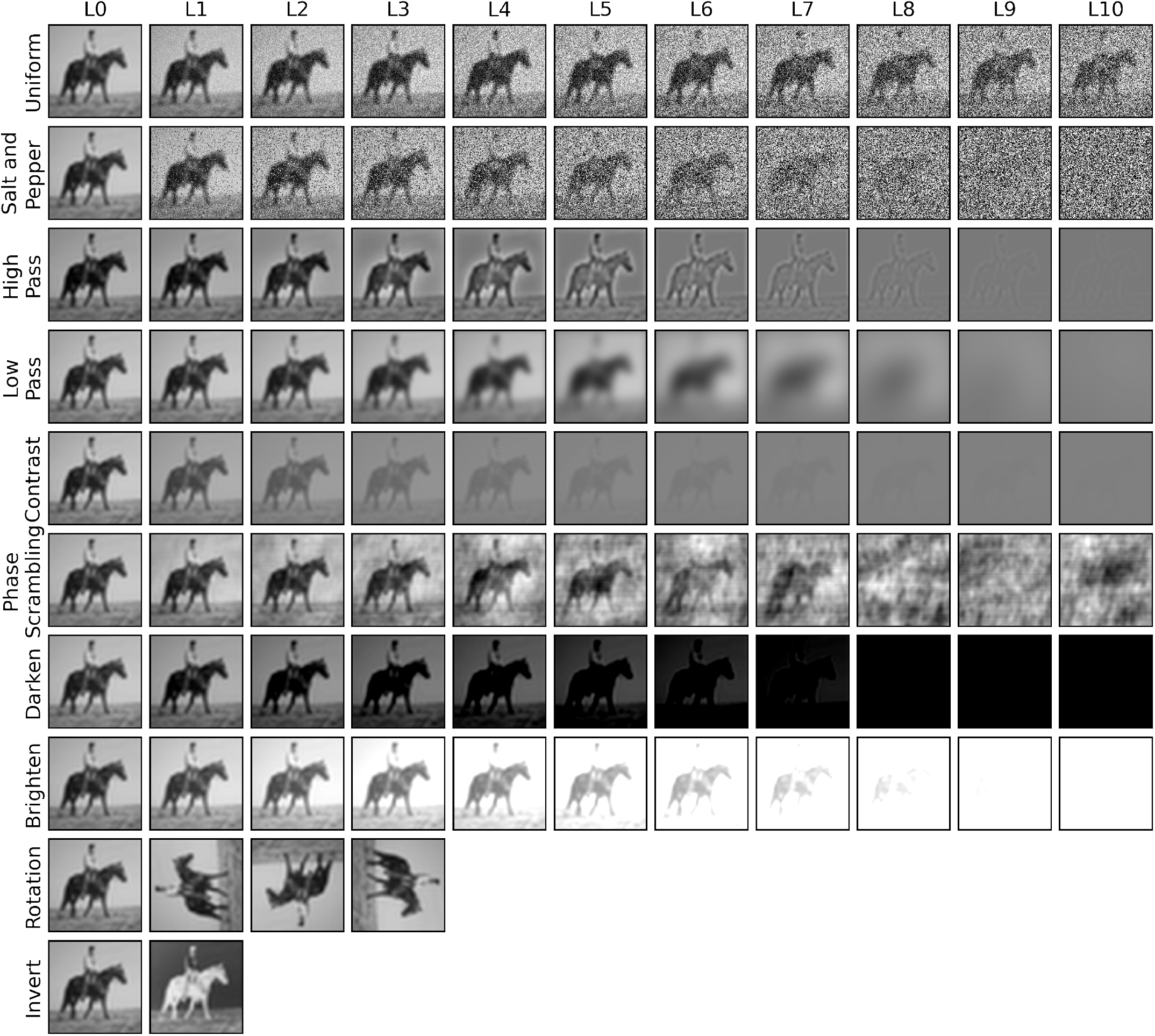
Noise perturbations at each level applied to an example `CIFAR-10` image.

#### 2.3.2 Generalisation Images

To test the networks’ abilities to classify images outside of the training set, we created a novel set of stylised (monochrome) test images (`CIFAR-10G`) for each of the ten `CIFAR-10` categories. These images contain mainly shape information, with very limited or no texture information at all, providing a means to assess a model’s ability to classify images without relying on the usual shortcut of spatially high-frequency information. Crucially these images are out-of-distribution (*o*.*o*.*d*.) in contrast to the reserved validation images which are independent and identically distributed (*i*.*i*.*d*.), as commonly used in machine learning research.

The images constituted three independent generalisation test sets: *line drawings, silhouettes* and *contours*. Each set had ten examples for each of the ten `CIFAR-10` categories. The contour images were derived from the silhouettes by hollowing out the shaded regions to leave only their outlines using the GNU Image Manipulation Program (GIMP). Finally, three additional sets were created by inverting the initial three sets. They came from a variety of internet sources but were all designated as free to use for commercial or other purposes. All six generalisation test sets are illustrated in Figure 3.

**Figure 3:**
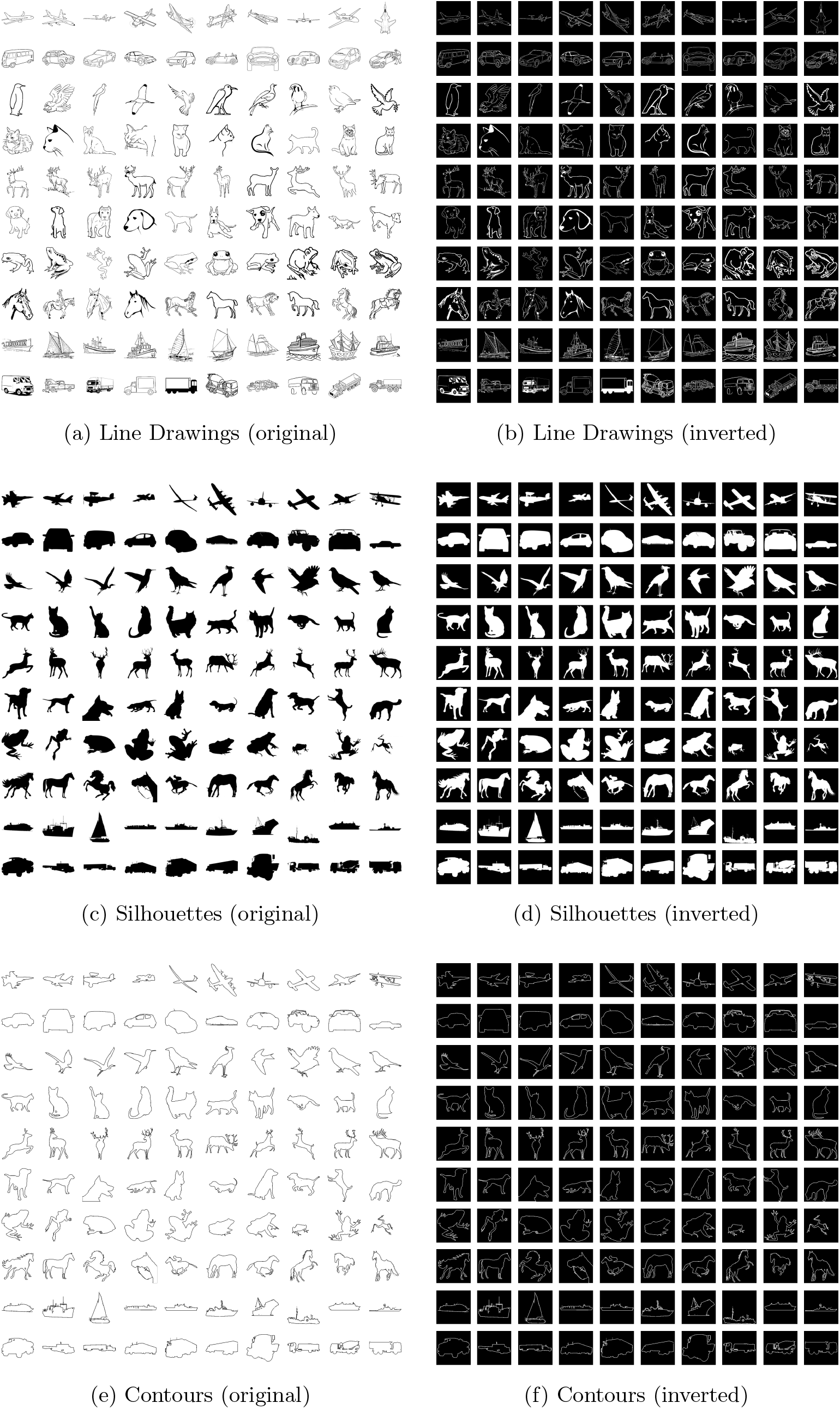
Generalisation test sets.

As a confirmation that these new generalisation image sets are truly *o*.*o*.*d*., the summary statistics (mean and standard deviation) of each image are plotted, along with those of the modified `CIFAR-10` train and validation sets, in Figure 4. Since the pixel intensities are rescaled to lie in the range [0, 1], the inverted images are reflected about the midpoint (*x* = 0.5) with respect to the original images they were derived from. While the training and validation sets lie on top of each other in the central region of the space, due to their sparse, largely binarised pixel intensities, the generalisation test sets lie on a manifold arcing around the edge of the space. This spatial separation demonstrates that they constitute out-of-domain test sets with respect to the `CIFAR-10` images.

**Figure 4:**
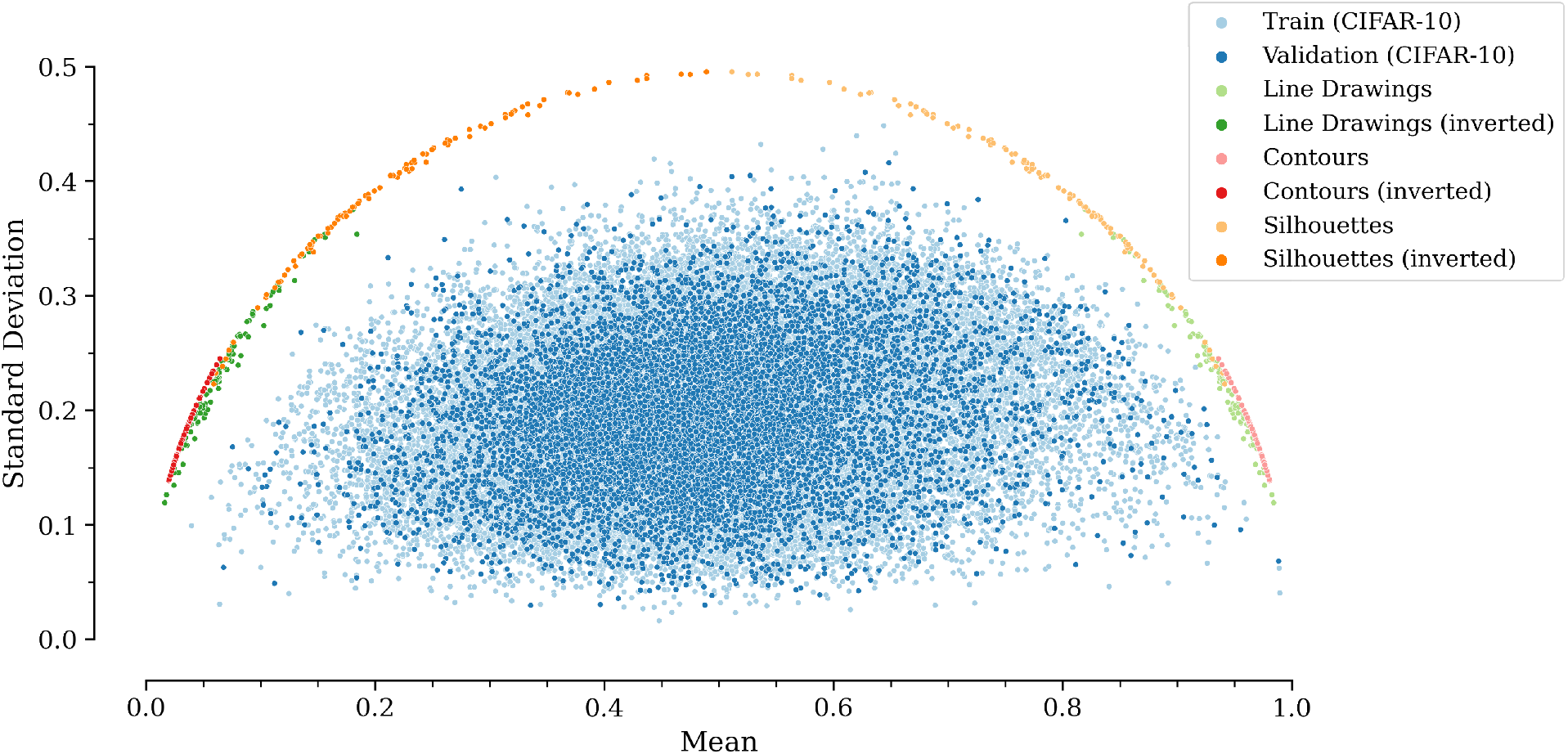
Distributions of image statistics. The `CIFAR-10` training and validation images are highly overlapping and occupy the central region of the space. Conversely, the generalisation images lie on a manifold forming an arc around this region, constituting out-of-domain test sets.

## 3 Results

### 3.1 Effect of the base model

We first checked that each model has broadly similar accuracy on the (unperturbed) `CIFAR-10` validation set, and that the pattern of differences due to the different convolutional “front-ends” holds for different “back-end” architectures. In Figure 5, the mean accuracy for each model (front-end / back-end combination) is plotted with the error bars representing the 95% confidence intervals calculated from five different random seeds.

**Figure 5:**
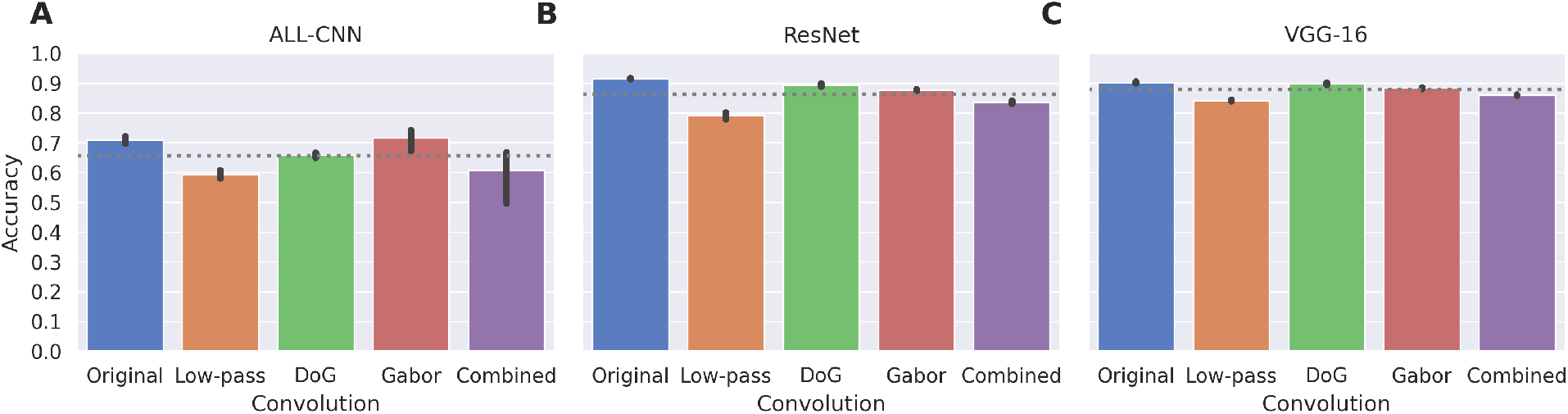
Classification accuracy on the `CIFAR-10` validation set. The `ResNet` and `VGG` models attained very similar levels of performance across all convolutional front-ends (around 90% accuracy) while `ALL-CNN` scored around 20% lower with more variability across front-ends. The line in each bar indicates the 95% confidence interval across the five random seeds. Grey dotted lines indicate the mean accuracy across convolutions for each base model architecture.

While the absolute levels of accuracy varied across the different architectures (with the performance of `ALL-CNN` being relatively low), importantly the relative pattern across front-ends remained very similar. We note from preliminary testing that, contrary to the trend of using deeper networks, the accuracy was largely unchanged after increasing the depth of the model from `VGG-16` **to** `VGG-19`. We note also that even the best performing models attain only around 90% accuracy, making them fall short from state-of-the-art for image classification. However, these figures serve as an adequate baseline for comparison to each model’s performance under more challenging and psychologically meaningful conditions.

### 3.2 Robustness to noise

After training to convergence on the modified (monochrome and upscaled) `CIFAR-10` training images, the networks were tested on the `CIFAR-10` validation set with various types and degrees of common noise perturbation, as described in Table 2.

Classification accuracy across the five runs of the `VGG-16` based models for each convolutional front-end under various levels of noise are given in Figure 6 as an example. The performance curves for `ALL-CNN` and `ResNet50` are given in Figures 12 and 13 respectively.

**Figure 6:**
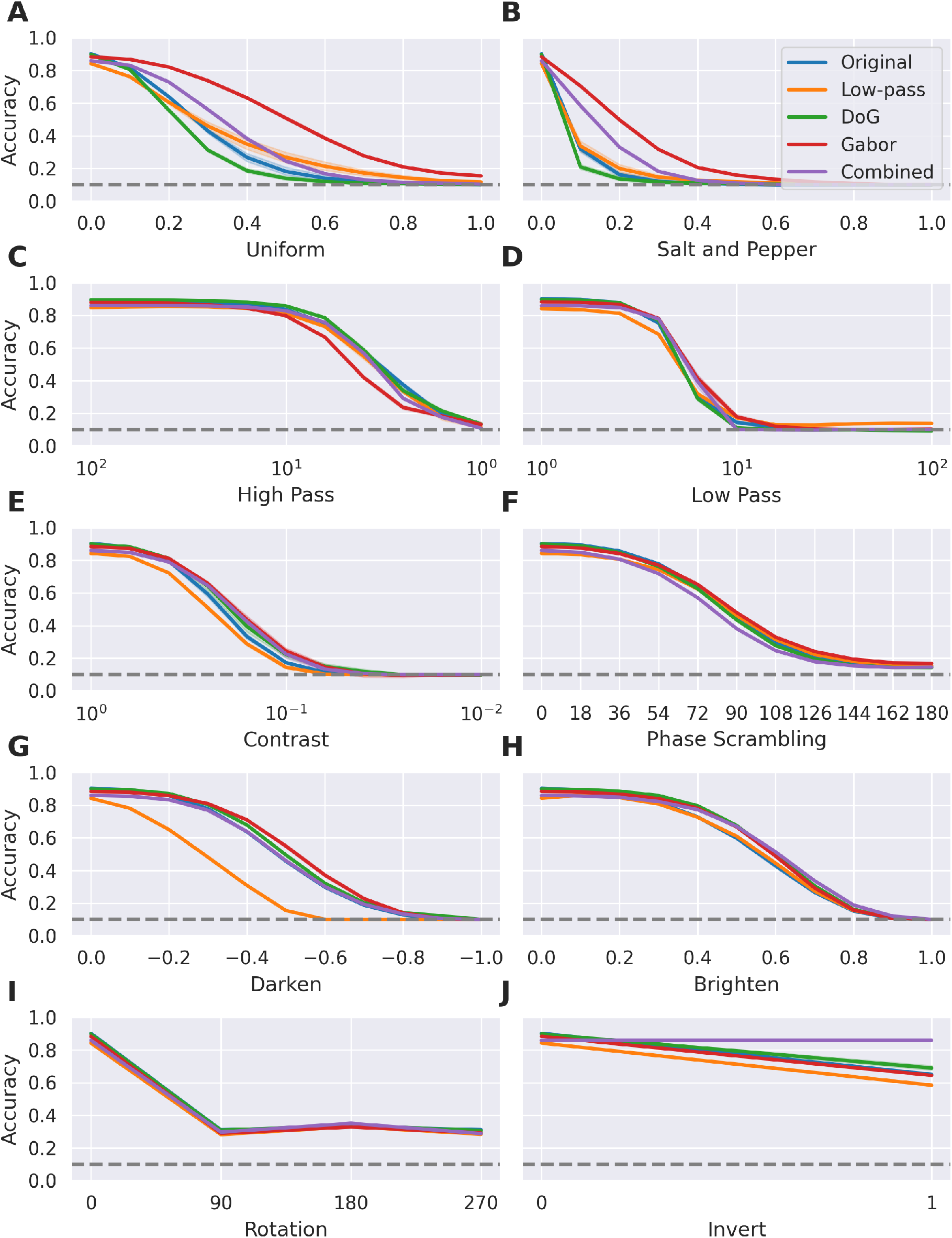
Classification accuracy of `VGG-16` based models under each type and degree of noise perturbation. The Gabor and Combined front-ends are particularly resilient to Uniform and Salt and Pepper noise, while the Combined front-end is able to recognise inverted images. Shading around each line indicates the 95% confidence interval across the five random seeds. The grey dashed lines represent chance level (10%) performance.

The perturbations used in this research are based upon previously published tests and common image degradations (Geirhos, Temme et al., 2020). As such, the fixed convolutional kernels used are not expected to lead to robustness in all cases. Earlier work suggests that resilience to uniform and salt-and-pepper noise should be improved (Malhotra et al., 2020; Malhotra et al., 2019). Additionally, biologically inspired filters are expected to be more resilient to brightened, darkened and reduced contrast images due to their regions of opponency which make them sensitive to spatial contrasts rather than absolute luminance levels. Conversely, end-to-end trained models are likely to maintain higher performance for high-pass filtered images owing to their preference for spatially high-frequency information such as their texture bias (Geirhos et al., 2019). For other perturbations such as rotation, we have no strong expectation of either an increase or decrease in robustness performance relative to the standard model.

In many cases, the biologically-inspired hard-coded convolutional front-ends (Gabor filters, Difference of Gaussians and Combined) are more or similarly robust to these types of image corruptions than their end-to-end trained counterparts (with the exception of High Pass perturbations). In particular, the Gabor and Combined models exhibited considerably more tolerance to Uniform and Salt and Pepper noise (Figure 6A&B) partly due to their smoothing effect. However, this characteristic alone can not entirely explain their large margin of improvement over other filters, due to the relatively poor performance of Low-pass filtered models under the same conditions, which serve as null models to test this idea. Instead, the combination of smoothing within a spatially structured kernel (i.e. elongated regions of opponency) appears to have helped reduce the effect of such unstructured noise on classification of natural images which consist of such spatially-structured features such as bars and edges (Bell & Sejnowski, 1997; Olshausen & Field, 1996). Gabor filters thereby offer the combination of edge-detection *and* spatial smoothing, helping them detect fundamental visual features while reducing their noise.

Interestingly, the Gabor-filtered networks tend to perform worse than the others when classifying images processed with High Pass filtering, (Figure 6C), presumably due to their bandwidth and spatial scale no longer being appropriate for the thinner edges and lines in this condition.

For perturbations such as phase scrambling and rotations (Figure 6F&I) all types of filter are quite similarly affected. Broadly comparable perturbation tolerance was also obtained for Contrast, Darken and Brighten (Figure 6E,G&H), with the exception of the Low-pass front-end, which was found to smooth away the finer details of the images, further reducing their contrast and reducing activation in subsequent layers.

While the Combined models exhibited similar patterns of tolerance to noise perturbation as the Gabor models, the absolute accuracy was typically lower. This may be explained by the information lost due to the extra DoG layer, as they may only occur in the visual system as a means of overcoming the retinal bottleneck (Lindsey et al., 2019). However, one notable exception is in the case of image inversion (Figure 6J) where most models drop by around 30% accuracy, whereas the Combined model is essentially unaffected. This is investigated further in Section 3.3.

While the original (unperturbed) validation images are *i*.*i*.*d*. with the training set (as illustrated in Figure 4), the models were not trained with any of the noise types, making this experiment a mild test of *o*.*o*.*d*. generalisation and a good test of more “real-world” image classification conditions.

### 3.3 Generalisation

As a strong test of *o*.*o*.*d*. generalisation, the models’ classification accuracy was assessed on the novel, stylised image test sets collected for this study (as shown in Figure 7). In almost all cases, networks with a Combined front-end scored highest, closely followed by Gabor models. One exception is on the silhouette test sets (original and inverted) where the Gabor front-end models outperformed the Combined models since these images had only edges (rather than other features such as lines) which the initial layer of DoG kernels are less sensitive to compared to Gabor kernels. Following those models, either the Difference of Gaussian or the Low-pass front-ends tended to slightly outperform the baseline Original models but were broadly comparable.

**Figure 7:**
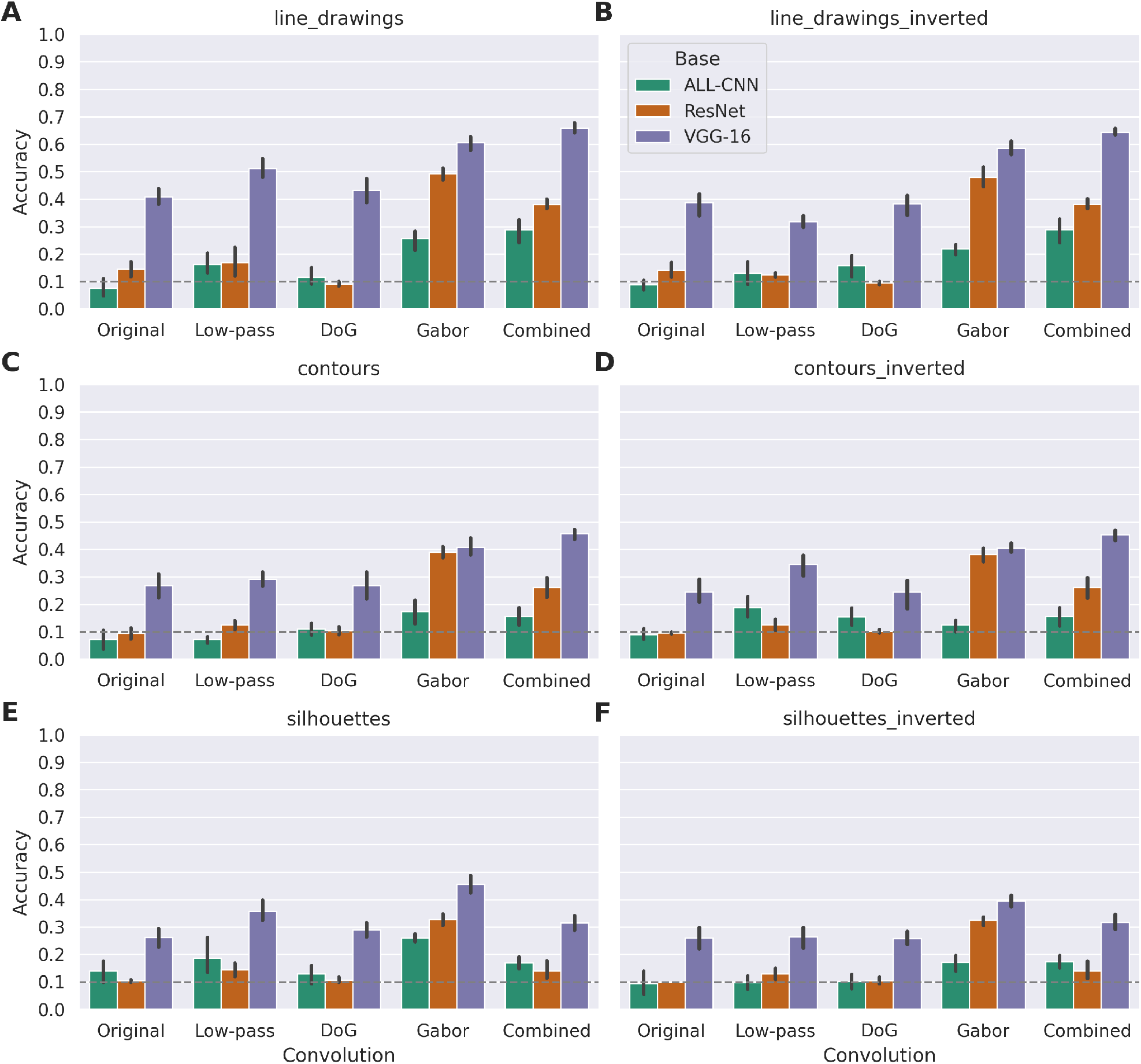
Classification accuracy for generalisation test sets. In classifying out-of-domain images, the Original (end-to-end) trained models typically score lowest. Classification accuracy with the Low-pass front end is slightly higher on average but less consistent across the test sets. The biologically-inspired convolutional front-ends have comparable performance (DoG front-ends) or substantially exceed the accuracy of the Original models (Gabor and Combined front-ends). Generally, all models score highest on the line drawings, with contours and silhouettes presenting the biggest challenges. The line in each bar indicates the 95% confidence interval across the five random seeds. The grey dashed lines represent chance level (10%) performance.

The original end-to-end trained models trail those which include a bank of Gabor filters (Gabor, Combined) by approximately 10% across generalisation test sets for `ALL-CNN`, 15 − 35% for `ResNet` models and 20 − 30% for `VGG` models. While there is clearly room for further improvement, these results demonstrate that a substantial margin in performance is conferred on standard DNNs in *o*.*o*.*d*. test images simply by fixing the form of the first layer of convolutions with biologically-plausible Gabor kernels.

Again the Combined front-end exhibits no performance drop associated with inverting the images, (see Figure 7, left column versus right column) unlike small but consistent drops for most other front-ends, especially the Low-pass models. Inspection of the activation patterns in the early layers of the Combined models reveals that the initial DoG layer provides an effective remapping of the inputs. Since for each DoG filter spatial scale and centre-surround ratio there is both an “on-” and “off-centre” receptive field, they can be matched to the inverted or original images (respectively) to yield the same activation pattern for each. Subsequently, the set of odd Gabor filters are then applied to these contrast-enhanced activation patterns to extract the edges as a foundation for more complex representations in subsequent layers (Figure 8). Essentially, having a layer of on- and off-centre DoGs followed by Gabor filters with equal and opposite phases means that opposite combinations of these filters could be matched to produce the same patterns of activation for both an original image and an inversion of it, as shown by the cosine similarity measures.

**Figure 8:**
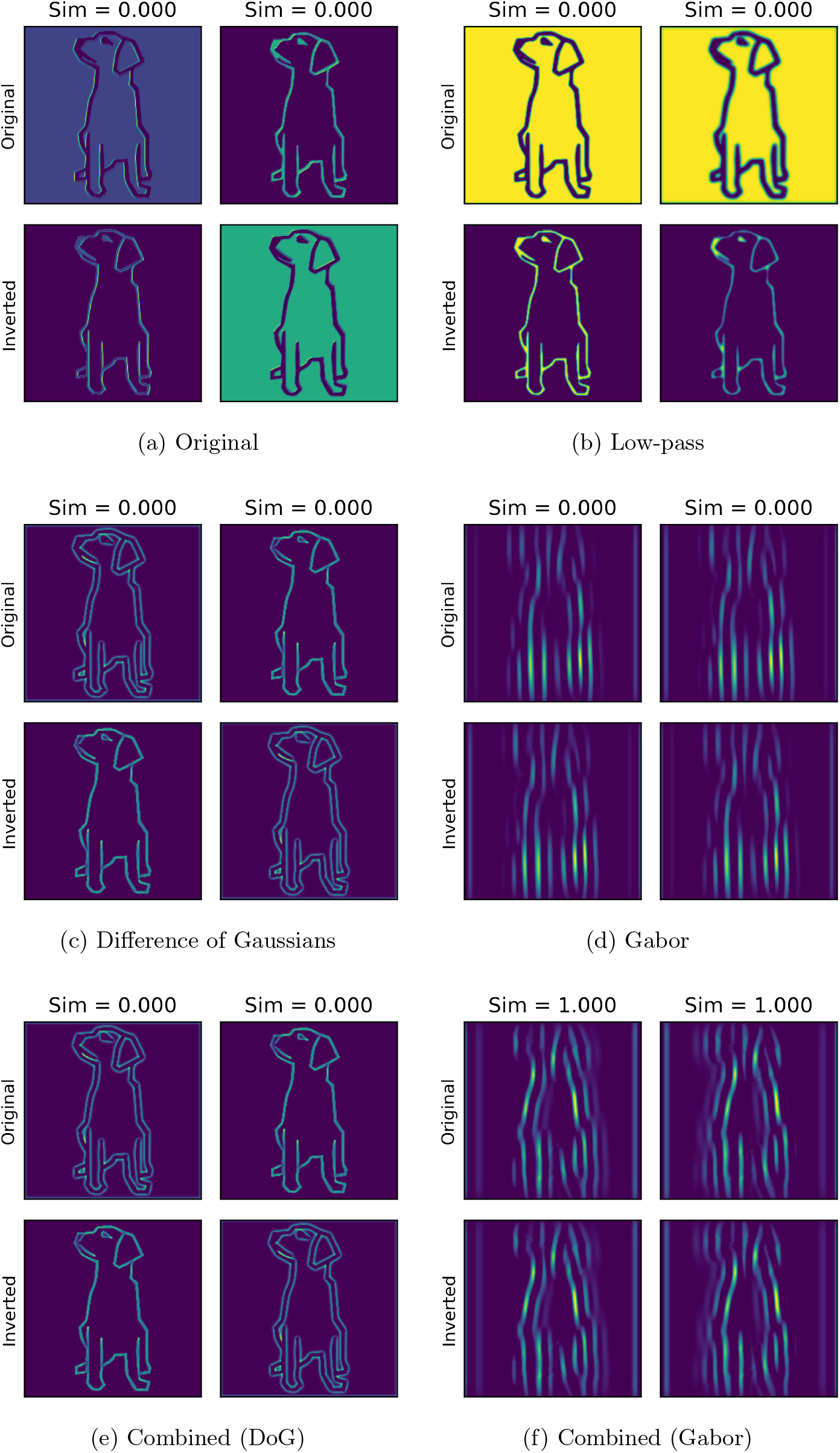
Activation maps generated from an example image and its inversion in the first four channels of the first convolutional layer(s). While the activations for the original and inverted images in the Gabor convolutions (d) appear similar to those in the Gabor layer of the Combined model (f), they are shifted with respect to one another. Conversely the preprocessing of the Combined front-end’s DoG layer (e) compensates for this phase-shift. The cosine similarities are shown for pairs of activations resulting from the original and inverted images.

In order to test how the convolutional front-ends affect the the models’ abilities to accurately classify *o*.*o*.*d* images under noisy conditions, we applied the same battery of image perturbations (shown in Figure 2) to the generalisation test image sets (shown in Figure 3). The results for one example generalisation test set — line drawings — are shown in Figure 9 on the `VGG-16` model architecture. Results for the other generalisation test sets (obtained with the same model architecture) are presented in the Appendix (Figures 14–18).

**Figure 9:**
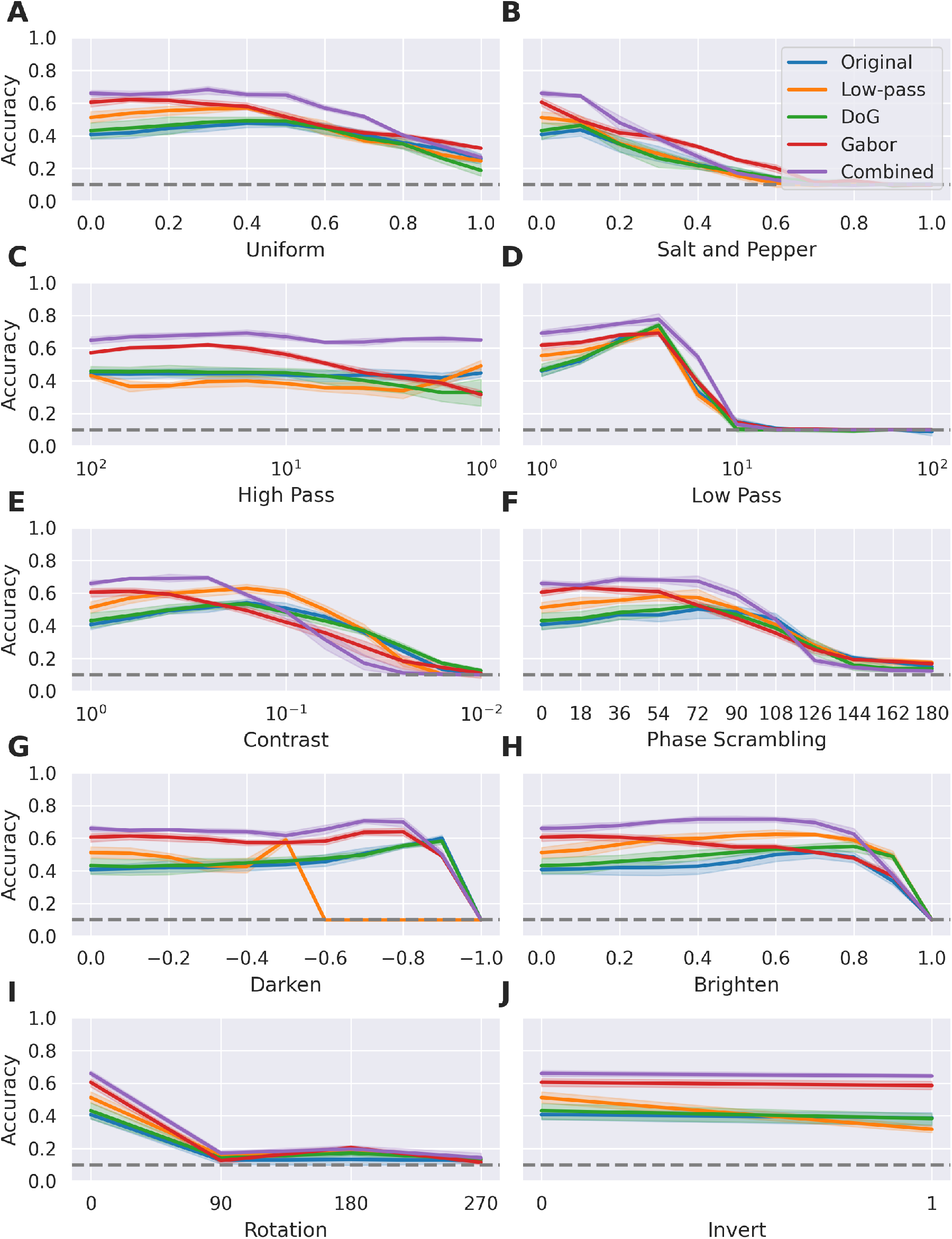
Classification accuracy of `VGG-16` based models on perturbed line drawings. The models with biologically-inspired convolutional front-ends (notably Gabor and Combined front-ends) typically maintain their advantage over the Original (end-to-end trained) models with the exception of low contrast and very dark perturbations. Shading around each line indicates the 95% confidence interval across the five random seeds. The grey dashed lines represent chance level (10%) performance.

Generally, the advantage conferred on the models by the biologically-inspired kernels holds across the set of perturbations, with their performance degrading more slowly relative to the Original front-end. In some cases the classification accuracy of the models improves slightly as the strength of perturbation increases. In the case of Low Pass perturbations (Figure 9D), this is likely due to the smoothing effect thickening the lines of the drawings, making them better able to to activate the filters of the first convolutional layer (whether fixed or learnt from naturalistic images). It is less clear why there would be an improvement to classification after applying, for example, Phase Scrambling but it is likely that such perturbations simply brought the images away from their outlying manifold (Figure 4) and closer to the image statistics of the training set.

### 3.4 Representations

In order to examine how the models’ internal representations are affected by the form of the initial convolutional kernels, the most activating features were determined for a selection of layers (Erhan et al., 2009). Initially, an image composed of random pixel intensities is presented to each model, which is then modified through gradient ascent for 1, 000 epochs to find the most activating feature(s) for that particular channel (subject to the random initialisation). The example channels were randomly chosen from the pooling layers, as they would effectively tile the preferred features of the preceding convolutional layer across the input canvas (although the convolutional layers produced very similar results). Representative examples of the most activating features for each of the `VGG-16` based models (for each front-end) are visualised in Figure 10.

**Figure 10:**
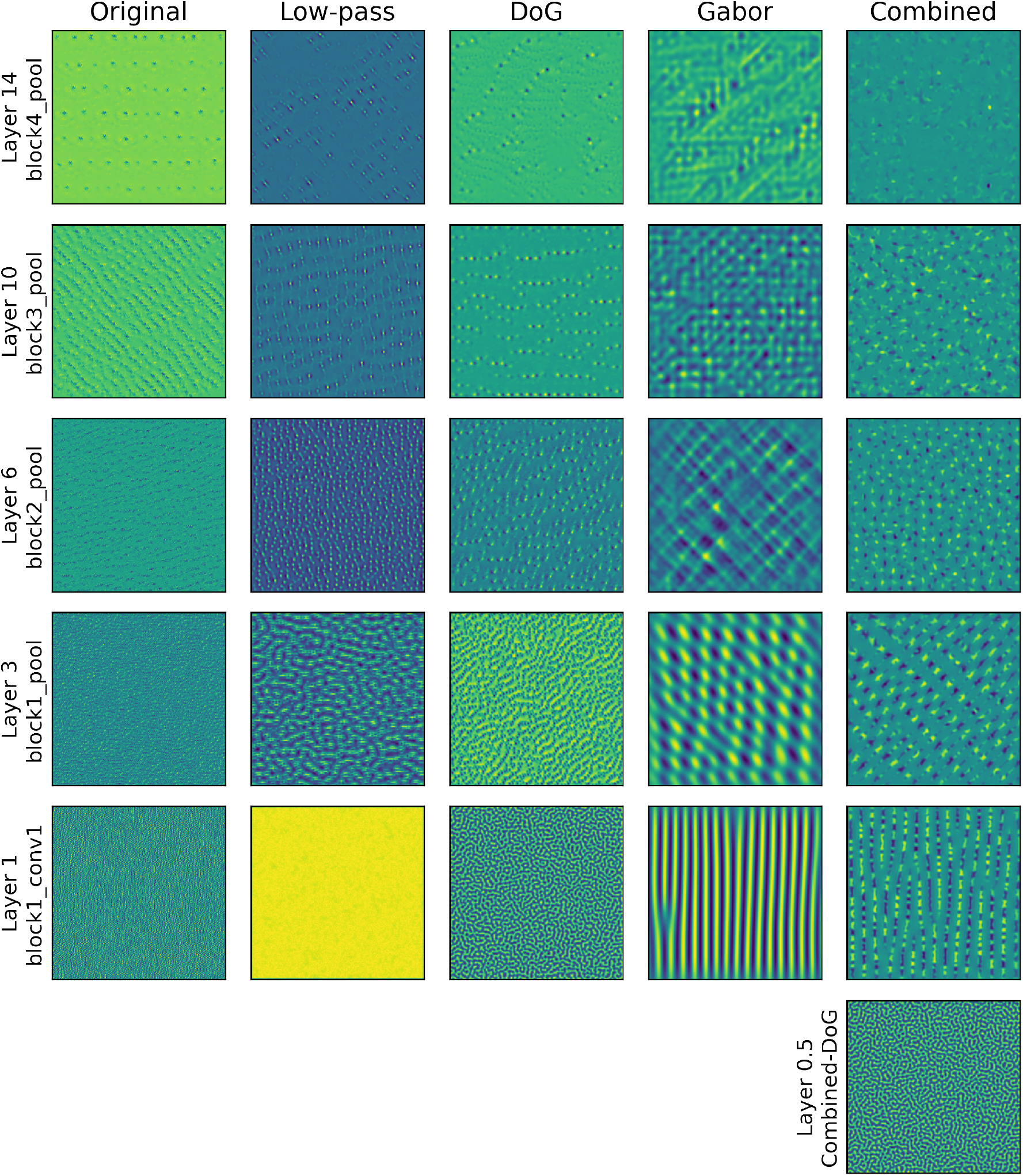
Most activating features for a selection of layers in models based on `VGG-16` with different initial convolutional layers.

There are clear differences in the most activating features across the different front-ends, evident in the visualisations, particularly in the earlier layers. The end-to-end trained (Original front-end) network prefers less structured and spatially very high-frequency patterns resembling noise. Conversely, the fixed kernel front-ends are all more activated by smoother, more structured patterns, with Turing patterns and oriented gratings observed for the Difference of Gaussians and Gabor front-ends respectively. It is often claimed that end-to-end training produces banks of Gabor-like units in DNNs that resemble simple cells of V1 (Krizhevsky et al., 2012). However, not only do these models learn a wide range of units, many of which do not resemble the receptive fields of neurons in early visual cortex, but our findings also highlight that hand-wiring the first convolutional layer(s) results in quite different learned representations in higher levels as well.

The learned features in the higher layers of the different models appear to be more similar than in early layers, in this case, appearing to converge to small blobs with antagonistic surrounds. Here it is hard to make any comparisons between the learned feature detectors in models and the brain because we have only a limited understanding of the features that drive single neurons in the higher levels of the visual system. Furthermore, any comparison between artificial and biological neural networks is further complicated by the fact that different methods of generating maximally activating images for single-units in ANNs can produce quite different outcomes, varying from unstructured noise to highly regular patterns, or even interpretable images (Nguyen et al., 2017). Similarly, different measures of single-unit selectivity provide very different estimates of selectivity (Gale et al., 2020). Importantly though, imposing fixed convolutional kernels in the early layers produces a major restructuring of the learned internal representations in otherwise standard DNNs — differences extending throughout the networks which are also found to have improved robustness and generalisation.

## 4 Discussion

The impressive performance of deep convolutional neural networks on various image classification benchmarks has led to a great deal of interest within the neuroscience community, where researchers are now exploring the similarity of human and DNN vision (Schrimpf et al., 2018). Indeed, optimising DNNs for image classification has been demonstrated to provide the best fit to observed neural activity in the primate visual system (Yamins et al., 2014) and yield similar patterns of representations across categories of objects as measured by Representational Similarity Analysis (Kriegeskorte, 2015). On this view, end-to-end training is the best approach to date for both image classification benchmarks *and* modelling human vision, so few inductive biases beyond convolution need to be incorporated.

However, here we show that hard-coding a filter-bank in standard DNNs that approximates the organisation of the early visual system improves the performance on noise-perturbed or out-of-domain images, compared to their standard (unconstrained) counterparts trained end-to-end. For example, the biologically constrained models were much better able to classify line drawings, mimicking humans infants, who can readily identify them without any explicit training (Hochberg & Brooks, 1962). Typical measures of model performance overlook many of these more interesting and elusive properties of biological visual perception, notably their ability to generalise, potentially driving research towards more narrowly defined goals and away from being more faithful models of vision.

It is also important to acknowledge that our biologically-inspired networks showed limited improvements compared to standard DNNs in some conditions, and in a few cases performed more poorly than their end-to-end trained counterparts. Clearly adding a fixed convolutional front-end is far from sufficient to overcome the limitations of current DNNs as models of human vision. This is perhaps not surprising, considering how different typical artificial and biological visual systems are, for instance the paradigm of rate-coding rather than temporal (spike) coding (Rullen & Thorpe, 2001), and the form of inputs they receive, such as static versus dynamic images. However, we argue that adding a biologically inspired front-end to standard DNNs represents a promising direction for advancement, especially for endowing them with better *o*.*o*.*d*. generalisation.

At an intuitive level, we may consider the benefit that biologically-inspired convolutional kernels confer on DNNs for classifying naturalistic images as arising from how they mimick the forms found through millions of years of evolution, which were useful and stable enough for decomposing natural visual scenes so as to be gradually enshrined in the genome. Both Difference of Gaussians and Gabor filters each incorporate antagonistic regions, whereby a feature of the visual scene (change in illumination) can be reliably detected and signalled in an energy efficient way (Vincent et al., 2005), the selection for which was likely driven by the need to constrain the high metabolic cost of transmitting information through spikes in the cortex (Lennie, 2003). By integrating their signals over a small spatial region, this also increases the reliability of the signal, by smoothing out sharp deviations in individually unreliable photoreceptors or pixels. In particular, developing Gabor-like receptive fields, with elongated, smoothed regions of opponency to signal changes, allows the organism or model to reliably detect bars and edges, which may then be used as the building blocks of shape — a key precursor to developing a concept of objects and a more reliable property for identification than low-level details such as texture.

Which features of biological vision need to be included in models in order to support human performance is still an open question. For example, the on- and off-centre receptive fields of retinal ganglion cells may simply be a means to compress the information from the photoreceptors through the retinal bottleneck in such a way as to be most faithfully reconstructed and expanded in the cortex (Vincent et al., 2005), without providing any additional benefit over Gabor-like receptive fields. This may explain the slightly mixed results with Gabor (only) versus Combined front-ends, such as their slightly weaker ability to classify silhouettes (compared to the Gabor front-end). If Gabor filters do indeed constitute an optimal “visual alphabet” as the first step in decomposing a natural visual scene when the information bottleneck is removed, then any additional (preceding) layer only serves to reduce the information content reaching them. It may be, however, that in order to cope with image inversions, additional pooling between Gabor filters of opposite phase is required — potentially an experimentally testable principle underpinning the finely structured organisation of the visual cortex.

The huge leap in performance and subsequent resurgence of interest in neural networks (then known as connectionist models) was brought about by the extraordinary increase in computational power through harnessing GPUs, along with access to much larger labelled image sets, which allowed much larger networks to be trained on vast amounts of training data (Krizhevsky et al., 2012). This trajectory still guides much of the community’s thinking on the best approach, typically eschewing such innate neurophysiological details and remaining largely empiricist in preferring end-to-end training. Despite a growing list of failures of such DNNs in classifying images under more challenging conditions (Geirhos et al., 2018; Geirhos, Temme et al., 2020), and demonstrations of striking differences between human and DNN vision (Dujmović et al., 2020; Malhotra et al., 2020), there is still the widespread view that many of these failures can be addressed by further improving the datasets that the models are trained on (Mehrer et al., 2017), or modifying the objective functions, including more emphasis on self-supervision (Chen et al., 2020) rather then constraining the models themselves with more inductive biases.

However, from examining the most activating features which are learnt throughout the networks, it is clear that constraining only the form of the initial convolutions has far-reaching effects for higher level representations which may impact the model’s ability to generalise. It is clear from this perspective, that even if benchmark-based summaries of the model’s performance are highly similar to those of their biological counterparts, it is unlikely that they are achieved in the same way, or that the same hierarchical organisation has necessarily developed (Thompson et al., 2021). It is only when testing models on more challenging datasets, that humans can readily identify, for example the distorted *i*.*i*.*d*. images or *o*.*o*.*d*. images of the present work, that these differences are manifest. The challenge in developing biological models of vision is to build models that explain or at least recapitulate core human visual capacities, such as scale and translation invariance (Blything et al., 2021; Han et al., 2020), the capacity to identify objects in novel orientations in 3D space (Erdogan & Jacobs, 2017) and tolerance to occlusion (Tromans et al., 2012), amongst many other human visual (limitations and) capacities.

Even when a bottleneck and other architectural constraints are added to networks to encourage the formation of (more) Gabor filters (Lindsey et al., 2019), there is still no hyper-column organisation of the filters or other potentially important details, and crucially, models still learn a wide range of other (spatially high-frequency) filters (Krizhevsky et al., 2012, Fig. 3), many of which do not occur in V1 or elsewhere as far as we know. This may help explain the brittleness of current DNNs with these extra kernels over-fitting to specific training sets, making the models less robust to distortions of *i*.*i*.*d*. images and considerably less able to recognise *o*.*o*.*d*. images. Ultimately, whether the V1 hyper-column structure is innately specified, or develops through (genetically guided) assimilation of early visual experience, current unconstrained DNNs trained end-to-end fail to capture the human ability to identify degraded images or generalise to out-of-distribution datasets.

### 4.1 Future work

To further enhance robustness and generalisation, it is likely that other modifications to the core components of ANNs are necessary, for example the addition of recurrent connections (Kietzmann, McClure et al., 2019; Kietzmann, Spoerer et al., 2019) or feedback connections (Kreiman & Serre, 2020). Also, in line with more standard approaches, it is undoubtedly important to also improve the training datasets and learning objectives in order to make models more similar to Infero-Temporal Cortex (IT), for example “soft” training labels (Peterson et al., 2019). In the current simulations we used supervised learning to train our models on `CIFAR-10`, and it would be interesting to see the impact of adopting different training objectives on larger datasets. For instance, there is some recent evidence that self-supervision on ImageNet can be used by networks to classify images more on the basis of shape compared to texture, consistent with the shape bias observed in humans (Geirhos, Narayanappa et al., 2020). In future work it will be important to understand how combining more inductive biases with better training regimes impacts on network performance.

### 4.2 Conclusions

In the presented work we have shown that adding biological filter banks to constrain standard DNN architectures reduces their capacity to find superficial solutions by “shortcut learning” (Geirhos, Jacobsen et al., 2020). In particular, our Gabor and Combined (DoG+Gabor) front-end models learned more structured internal representations, were more robust to a number of common noise perturbations, and most importantly, showed better generalisation to our novel *o*.*o*.*d*. test sets. We take these findings as evidence that researchers should incorporate more biological constraints in DNNs to better mimic human performance, and indeed, it may be an important step in developing machine learning systems that generalise better. More generally, we also advocate a wider perspective on model evaluation than a narrow focus on common benchmark scores, as this is likely to lead to models which miss many of the more interesting and useful properties of human vision.

## Acknowledgements

Funding: This project has received funding from the European Research Council (ERC) under the European Union’s Horizon 2020 research and innovation programme (Grant Agreement No. 741134).

We also thank Alex Hernandez-Garcia for his implementation of `ALL-CNN`.

## Appendix

**Figure 11:**
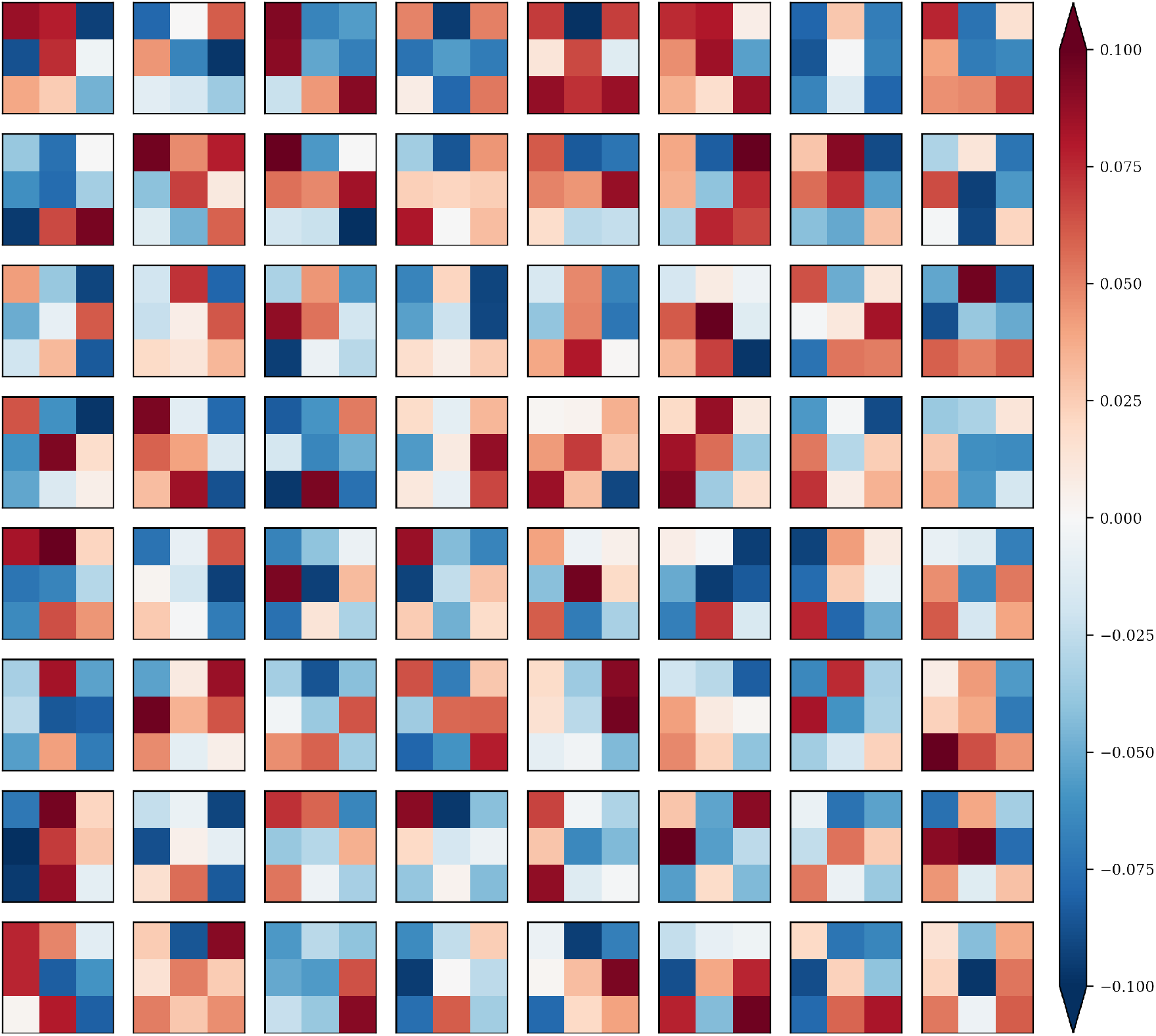
Illustration of the first convolutional layer kernels obtained through end-to-end training in the `VGG-16` model architecture.

**Figure 12:**
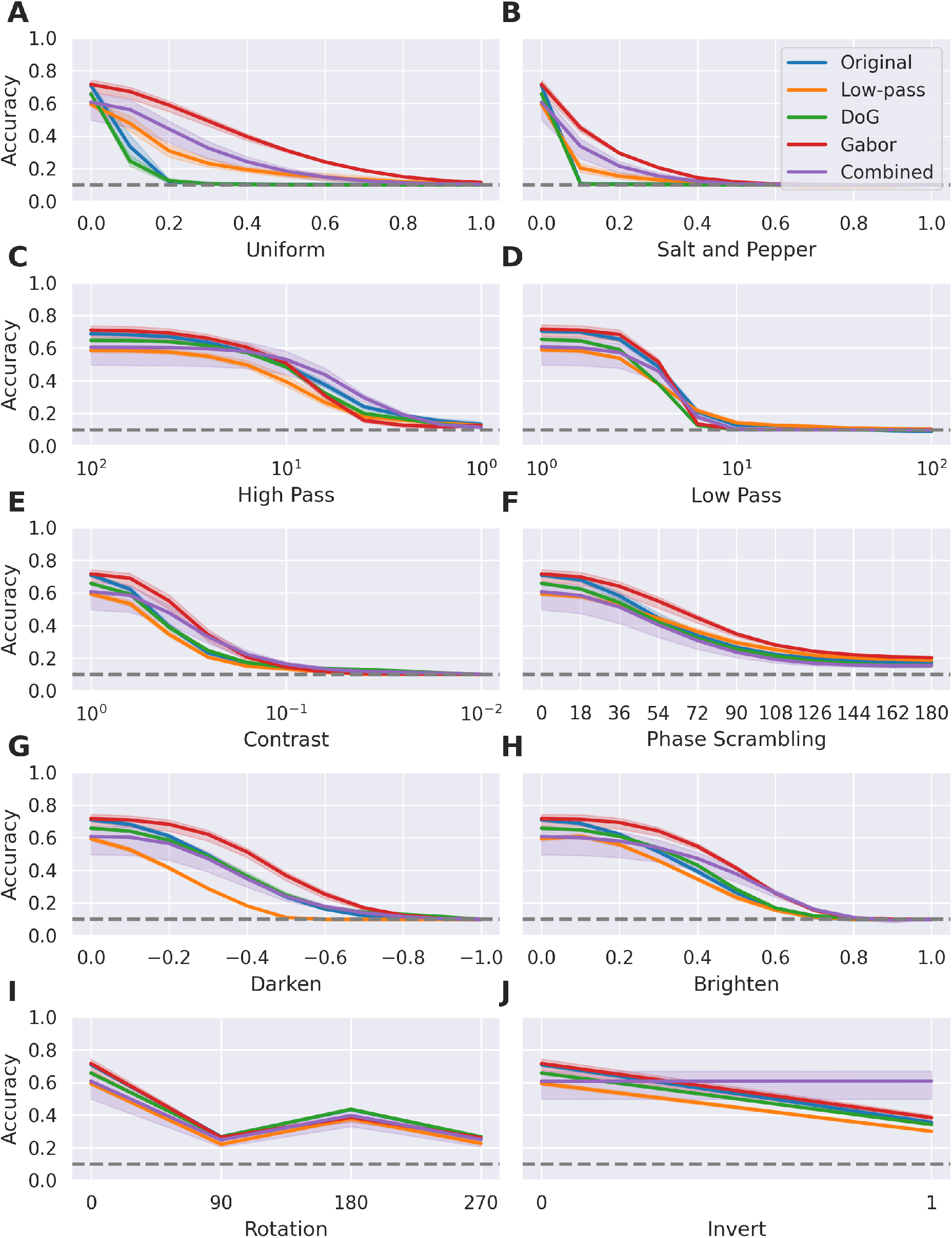
Classification accuracy under different types and degrees of noise perturbation for `ALL-CNN` based models. Shading around each line indicates the 95% confidence interval across the five random seeds. The grey dashed lines represent chance level (10%) performance.

**Figure 13:**
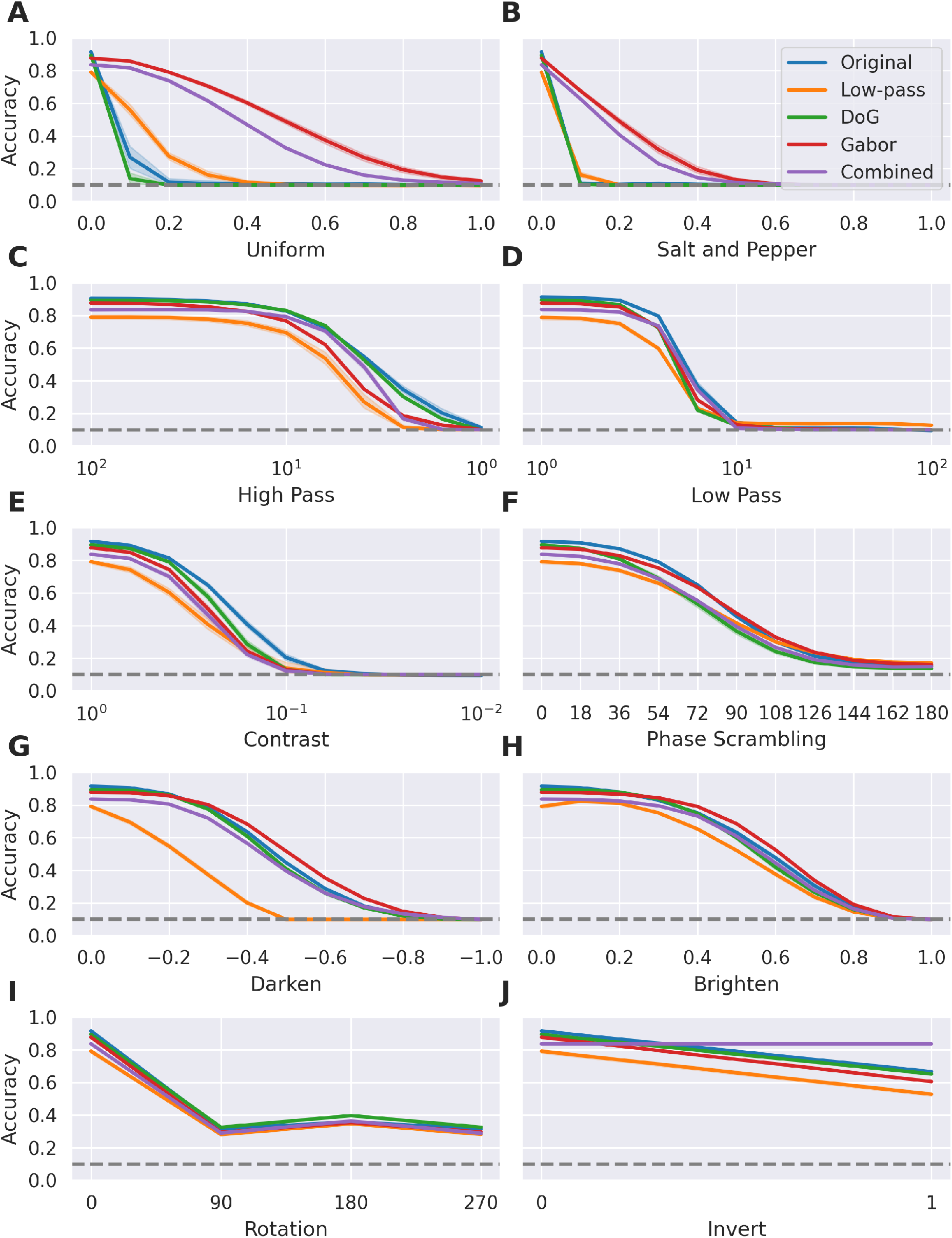
Classification accuracy under different types and degrees of noise perturbation for `ResNet50` based models. Shading around each line indicates the 95% confidence interval across the five random seeds. The grey dashed lines represent chance level (10%) performance.

**Figure 14:**
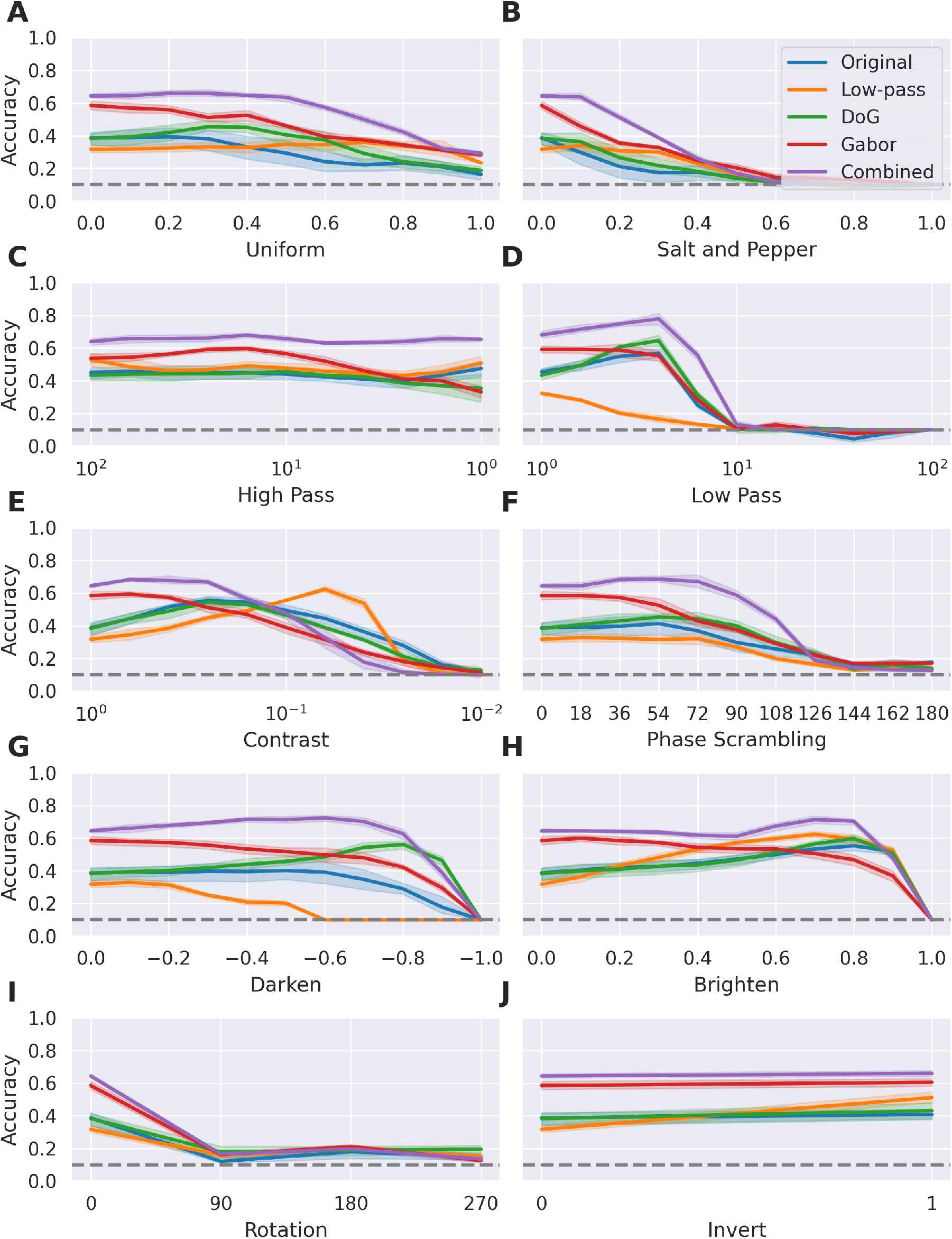
Classification accuracy of `VGG-16` based models on perturbed inverted line drawings. The models with biologically-inspired convolutional front-ends (notably Gabor and Combined front-ends) typically maintain their advantage over the Original (end-to-end trained) models with the exception of low contrast perturbations. Shading around each line indicates the 95% confidence interval across the five random seeds. The grey dashed lines represent chance level (10%) performance.

**Figure 15:**
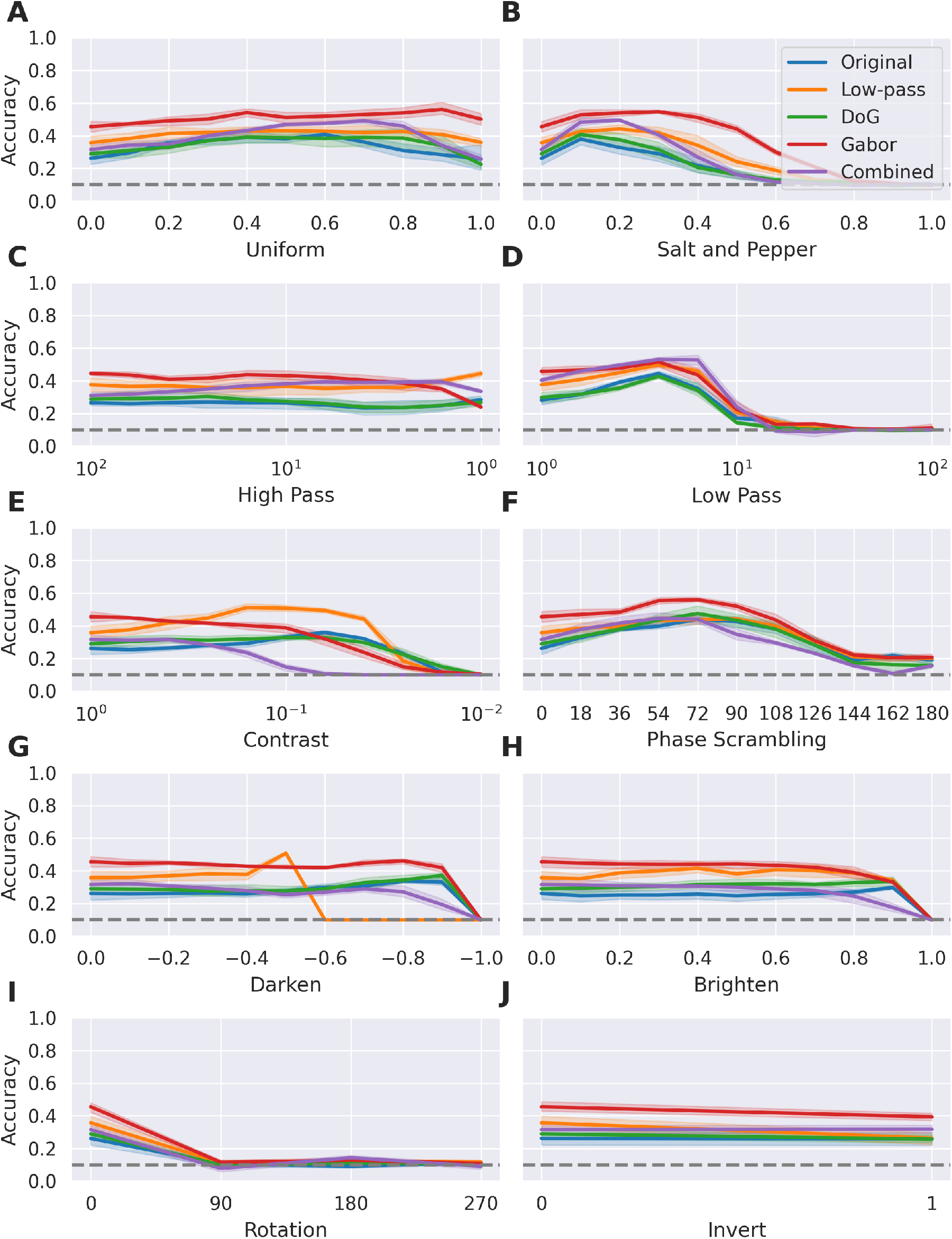
Classification accuracy of `VGG-16` based models on perturbed silhouettes. The models with biologically-inspired convolutional front-ends (notably Gabor and Combined front-ends) typically maintain their advantage over the Original (end-to-end trained) models with the exception of low contrast perturbations. Shading around each line indicates the 95% confidence interval across the five random seeds. The grey dashed lines represent chance level (10%) performance.

**Figure 16:**
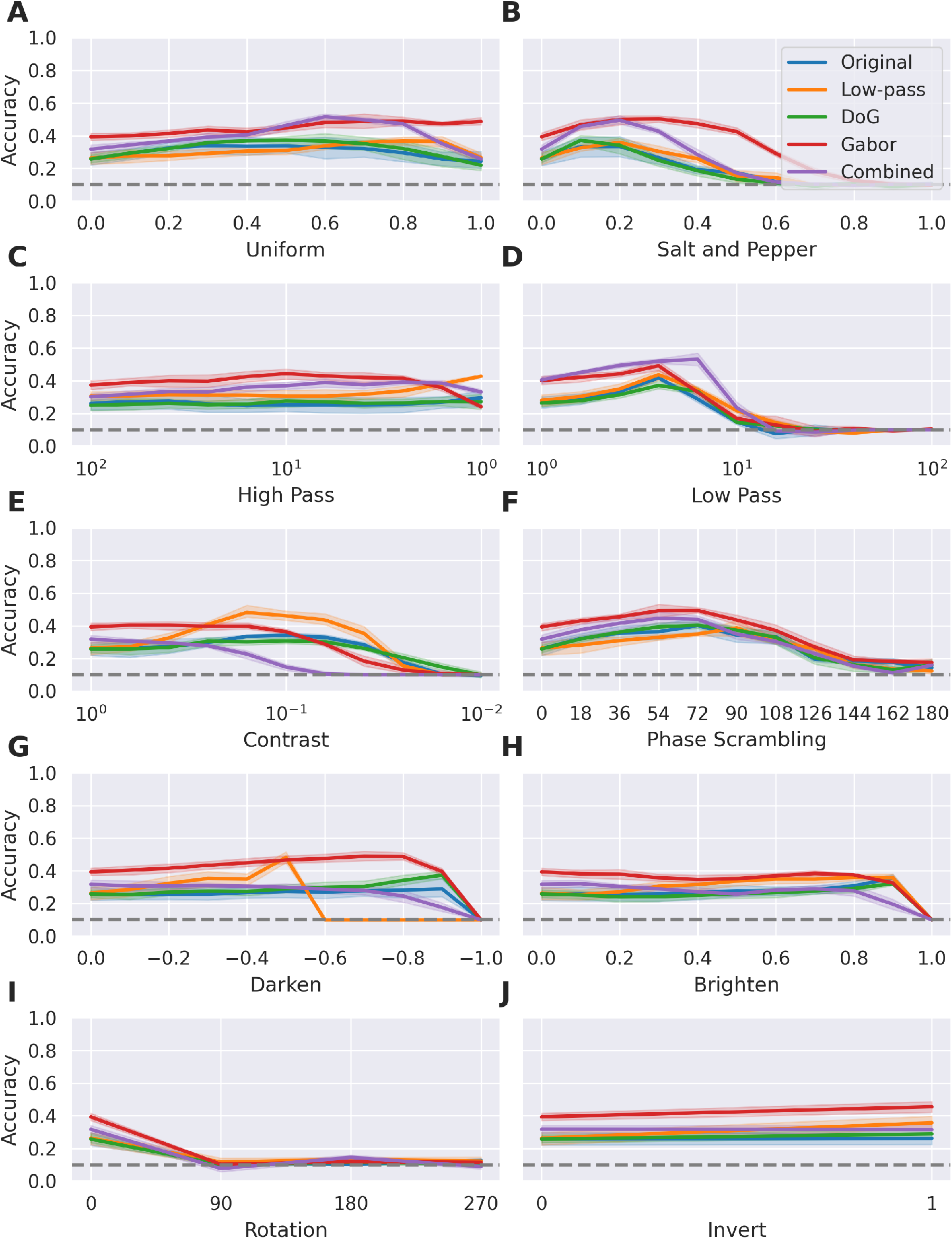
Classification accuracy of `VGG-16` based models on perturbed inverted silhouettes. The models with biologically-inspired convolutional front-ends (notably Gabor and Combined front-ends) typically maintain their advantage over the Original (end-to-end trained) models with the exception of low contrast perturbations. Shading around each line indicates the 95% confidence interval across the five random seeds. The grey dashed lines represent chance level (10%) performance.

**Figure 17:**
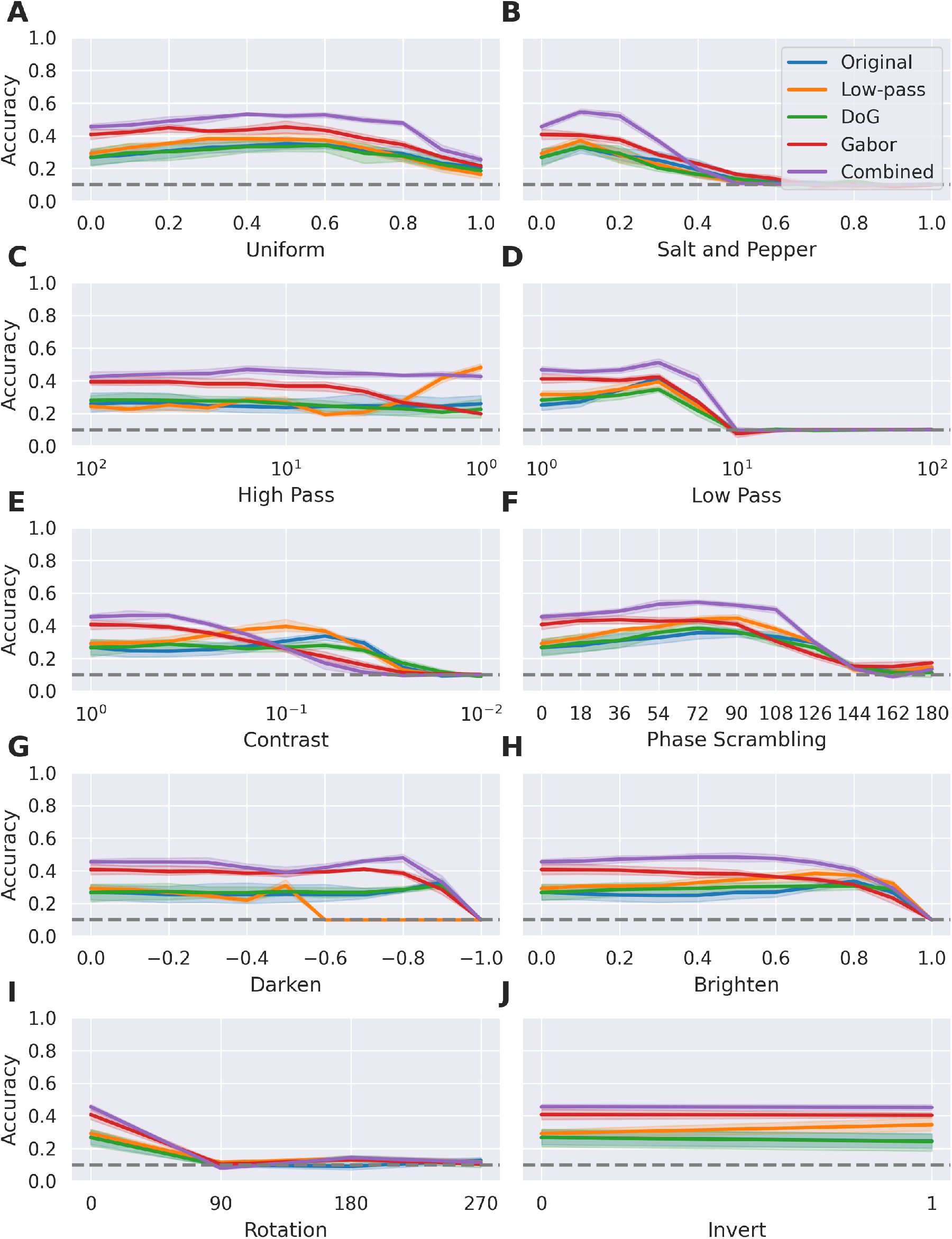
Classification accuracy of `VGG-16` based models on perturbed contours. The models with biologically-inspired convolutional front-ends (notably Gabor and Combined front-ends) typically maintain their advantage over the Original (end-to-end trained) models with the exception of low contrast perturbations. Shading around each line indicates the 95% confidence interval across the five random seeds. The grey dashed lines represent chance level (10%) performance.

**Figure 18:**
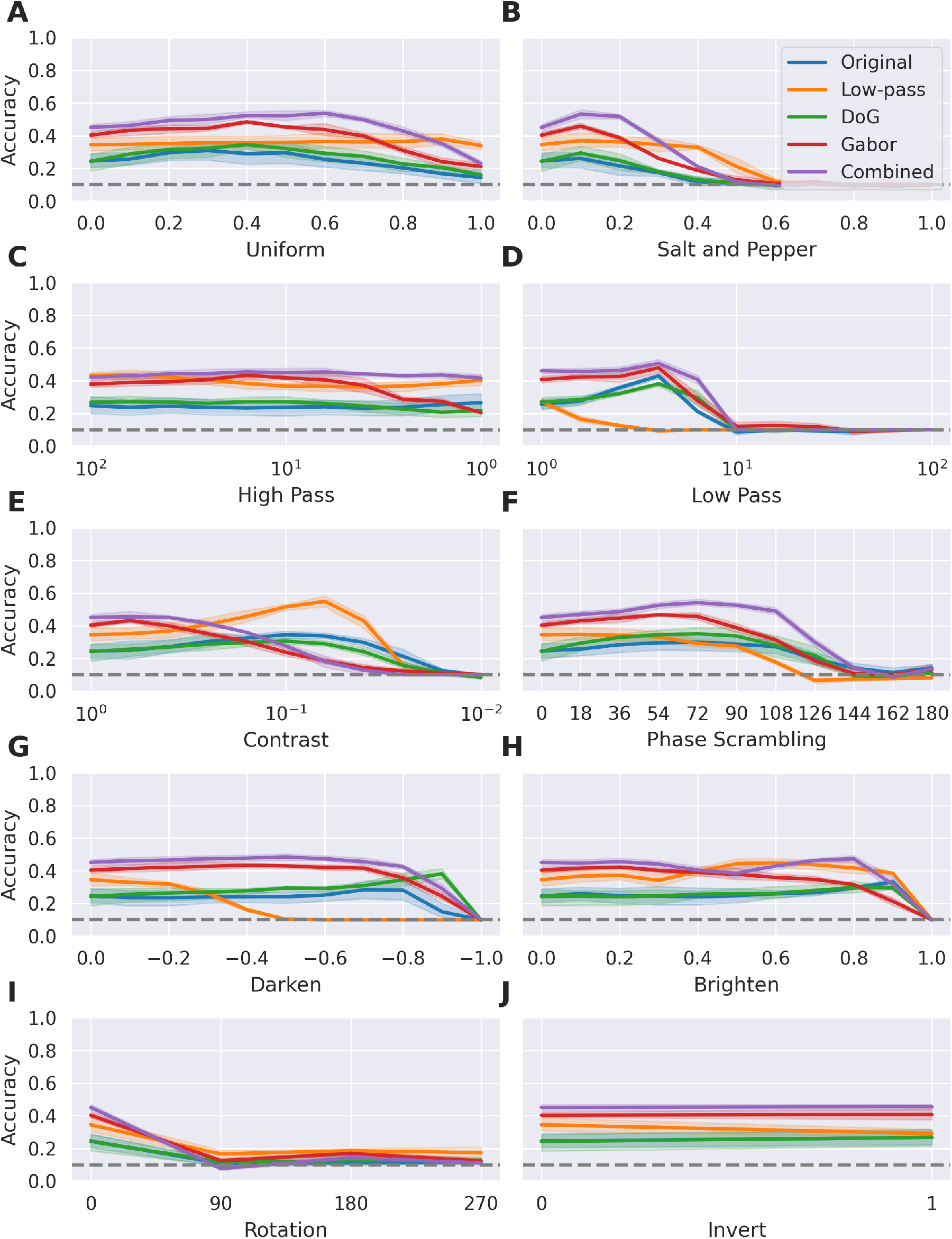
Classification accuracy of `VGG-16` based models on perturbed inverted contours. The models with biologically-inspired convolutional front-ends (notably Gabor and Combined front-ends) typically maintain their advantage over the Original (end-to-end trained) models with the exception of low contrast perturbations. Shading around each line indicates the 95% confidence interval across the five random seeds. The grey dashed lines represent chance level (10%) performance.

## Notes

https://github.com/bdevans/BioNet

### Competing Interest Statement

The authors have declared no competing interest.

### Summary of Updates

Results for VGG-19 were replaced with ResNet50 (Revised Figs 5, 6 & 7); Figures 1&11 were added to illustrate the convolutional kernels; The previous Figure 6 was removed; New results were added in Figures 9, 14--18, examining tolerance to perturbations in with the generalisation test images; Associated minor revisions to the text were made, including revising Equation 5, adding new references and extending the discussion.

